# Principles of neocortical organisation and behaviour in primates

**DOI:** 10.1101/2025.07.17.665410

**Authors:** Katja Heuer, Nicolas Traut, Leandro Aristide, S. Faezeh Alavi, Marc Herbin, Rogier B. Mars, Rajeev Mylapalli, Shaghayegh Najafipashaki, Tomoko Sakai, Mathieu Santin, Víctor Borrell, Roberto Toro

## Abstract

**One sentence summary:** The study of brain MRI from 70 different primate species reveals a fundamental principle of neocortical organisation and behaviour driven by mechanical morphogenesis.

The development and evolution of neocortical organisation is typically explained by the interaction of two fundamental factors: genetics and experience-dependent processes. Morphogens and signalling molecules would orchestrate the formation of neocortical areas and connection networks, which are later refined through exposure to environmental stimuli. Evolutionary changes to these genetic programs are thought to account for the diversity of brains and behaviours observed in extant species. However, our phylogenetic comparative study of primate neuroanatomy and behaviour shows this view is incomplete. Using brain MRI from 70 primate species we observed that not only the degree of folding but also the folding pattern changes continuously with brain volume, independently of phylogenetic position. To better understand the consequences of this continuity we focused on New and Old World monkeys which diverged approximately 47 million years ago. Large New World monkeys, such as capuchins, have a significantly larger and more folded neocortex than many of their close phylogenetic relatives, whose brains are barely folded. Notably, in addition to folding, their thickness and connectivity patterns were almost identical to those of phylogenetically distant Old World monkeys. Combined analyses of MRI and endocasts from 105 primate species indicated that the highly folded neocortex of large New World monkeys evolved independently from a common ancestor with a small, unfolded brain. Remarkably, across all 70 species, behavioural similarity correlated substantially more with neuroanatomical similarity than with phylogenetic similarity. Our results challenge the prevailing explanation of the development and evolution of neocortical organisation. We propose that the “capuchin anomaly” can be resolved by incorporating mechanical morphogenesis, alongside genetics and experience, as a third fundamental factor. Growth-driven mechanical instabilities would produce similar neuroanatomical organisation patterns and behaviours, emerging independently of the specific genetic determinants of that growth.

## Main

The neocortex is a unique mammalian innovation and the largest part of the human brain, linked to complex cognitive functions such as planning and language. Neocortical organisation is characterised by the strong regularity of its radial dimension: neuronal cell bodies form a distinctive 6-layer motif with dendritic trees and intra-cortical axons creating a dense plexus of synaptic connections. Whereas the thickness of the neocortex is remarkably stable – 1.5 mm in mice and 2.5 mm in humans – its tangential extension spans over 4 orders of magnitude. Variations in the 6-layer motif, their cellular composition and connectivity patterns, allow us to identify specialised neocortical areas. The number of these areas increases linearly with brain size, and the number of neural connections that link them increases exponentially. Large neocortices are also folded, with folding patterns being characteristic of each species and related to their areal, connective and functional organisation (Brodman 1909, Fischl et al. 2007).

The prevailing paradigm is that neocortical organisation is established during early development by two fundamental factors: genetics and experience (Rakic 1988, O’Leary 1989, Krubitzer 2007, Imam and Finlay 2020). Morphogens and signalling molecules lay down a coordinate system over which patterned gene expression determines the formation of neocortical regions and their basic connectivity. By differentially regulating various aspects of cell proliferation, migration and maturation, these genetically encoded patterns also determine the species-characteristic pattern of neocortical folding (Del Valle Anton and Borrell 2022, Akula et al. 2023).

Accordingly, if differences in folding patterns, neocortical arealisation, connectivity and behaviour observed across species reflected their different genetic makeup, then closely related species should exhibit a more similar neocortical organisation and behaviour than those farther apart. However, our phylogenetic comparative analyses of the neuroanatomy and behaviour of 70 primate species suggest that this view is incomplete.

We show that across primate brains, from the small and unfolded to the large and profusely folded, folding patterns are strongly related to brain volume, independent of phylogenetic position: an evolutionary trajectory built by combining brains of increasing volume shows a striking continuity, despite jumping across far apart branches in the phylogenetic tree.

To better understand the consequences of this continuity, we focused on New World and Old World monkeys, which provide a unique “natural experiment”. Primates appeared about 74 million years ago. From Africa, the common ancestor of New World and Old World monkeys crossed the Atlantic Ocean and dispersed into South America some 47 million years ago – about the same time when the common ancestor of whales and hippopotamuses returned to the ocean. After an early burst of speciation 21 million years ago, the ancestral New World monkeys evolved into a diverse group of primates, spanning the range from small marmosets and tamarins to the larger capuchins and sapajus. Our analyses indicate that the ancestral New World monkey had a small, barely folded brain. From there, large New World monkeys have expanded their brains at a pace only paralleled by humans, reaching a volume comparable to that of Old World monkeys such as macaques.

Our results show that in addition to the pattern of cortical folding, the patterns of cortical thickness and whole-brain connectivity, and even the behaviour of capuchins, are much more similar to those of Old World monkeys with similar brain size than to those of phylogenetically closer New World monkeys with a smaller brain. This “capuchin anomaly” challenges the current paradigm, as it would require a parallel evolution of the genetic encodings of areal, connective and folding patterns, which would not leave traces at the genomic level. We propose that this anomaly is resolved by conceiving a fundamentally different kind of convergence – one driven by mechanical constraints rather than genetic adaptation. Mechanical morphogenetic processes act as a third fundamental factor, alongside genetic and experience-dependent factors, in determining neocortical organisation and behaviour. Overall, our findings reveal a mechanically driven form of convergent evolution: when brains reach a certain volume, similar neuroanatomical patterns emerge due to shared mechanical constraints. This leads not only to convergent folding patterns but also to convergent cortical thickness maps, whole-brain connectivity, and even behaviour. Crucially, this process is independent of the genetic mechanisms responsible for brain volume expansion, which are known to be highly polygenic.

### The folding pattern of primates is strongly determined by brain volume, irrespective of phylogenetic position

The correlation between brain volume and degree of folding across mammals is well established (Prothero and Sundsten, 1984). We reconstructed the cortical surfaces of 70 primate species (Fig. 1), where this correlation was very strong (r=0.95, CI=[0.92, 0.97] see also Rogers et al. 2010, Heuer et al. 2019). Using surface morphing, we observed that not only the degree but also the pattern of primate brain folding was strongly related to brain volume. Video 1 shows the evolutionary trajectory of neocortical expansion across 54 primate species, including all major groups of primates, from small lemurs to great apes. Ordered solely by brain size – ignoring phylogenetic relationships and jumping across far apart branches in the phylogenetic tree – a strikingly continuous and predictable trajectory of folding pattern changes emerges, illustrating a common underlying organising principle (video available at https://doi.org/10.5281/zenodo.16033813).

**Figure 1.**
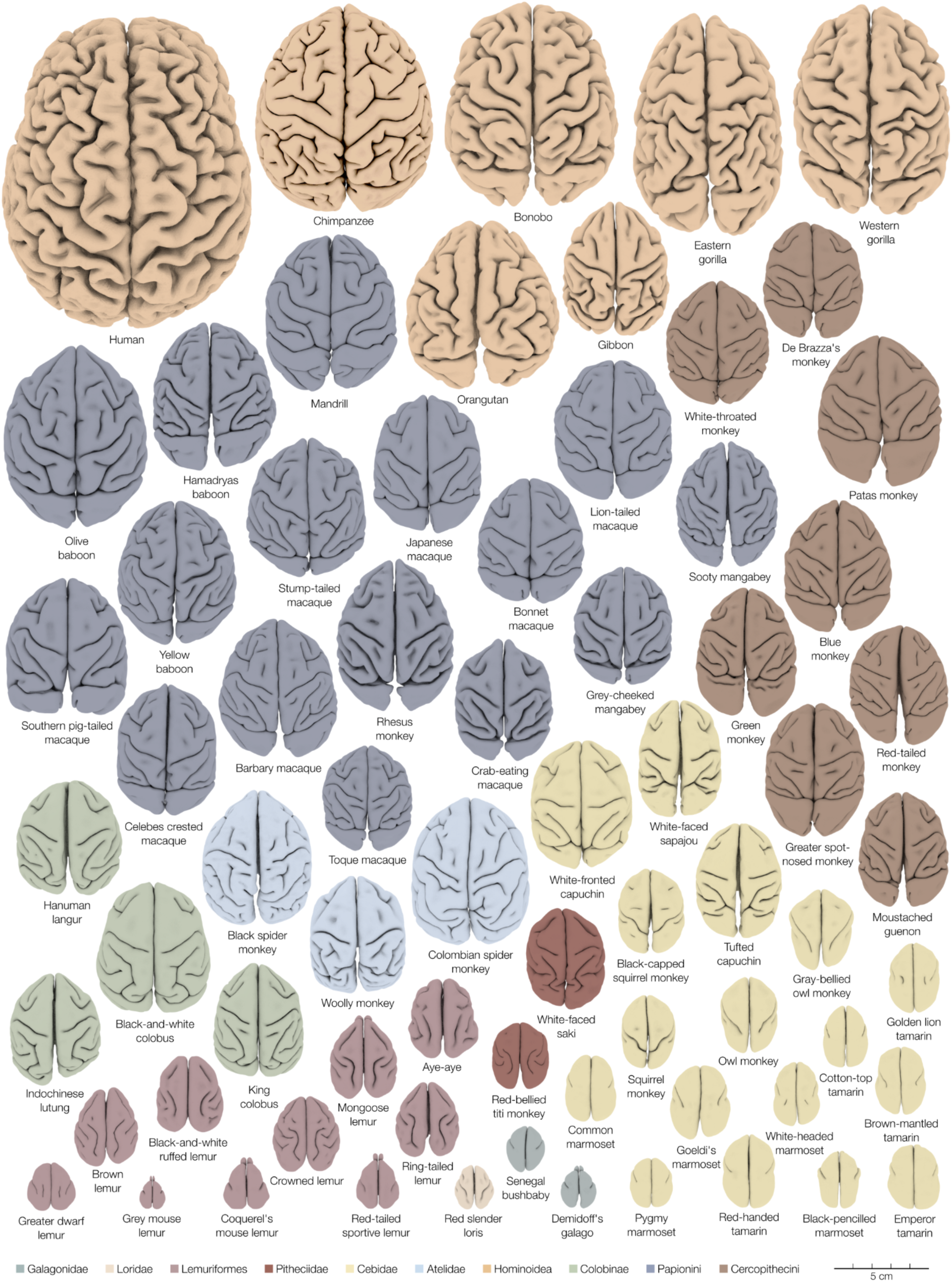
Dorsal view of the reconstructed cerebral hemispheres of 70 primate species. Brain surfaces include species from all major groups of primates. Colours correspond to the different families and indicate phylogenetic relationship, and brains are represented from largest on top to smallest at the bottom. The scale is the same for all brains. High-resolution version: https://doi.org/10.5281/zenodo.15969765

### The common ancestor of New World and Old World monkeys had a small, nearly smooth brain

To better understand the consequences of the continuous relationship between folding pattern and brain volume, we focused on New World and Old World monkeys, where species have evolved similar brain volume independently. We estimated the neuroanatomy of their common ancestor using phylogenetic comparative methods with two complementary types of data: surface reconstructions from brain MRI data and brain endocasts. Both approaches led to the same conclusion: large New World monkeys evolved from an ancestor with a small, nearly smooth brain. First, we measured total volume, total surface area, gyrification index, number of folds, total length of folding, average fold width and average fold depth in brain surfaces of 70 primate species. The phylogenetic tree for our species is illustrated in Fig. 2, along with several additional examples of brains from diverse branches that have a similar volume, and pairs of brains from closely related species with dissimilar volumes. Among different evolutionary models, the Brownian Motion model provided the best fit (in this model, phenotypic difference between species varies proportionally to phylogenetic branch length separating them from their common ancestor). Using this model, we estimated the evolution of total cerebral volume, surface area, and folding along the phylogenetic tree, shown in the phenograms in Fig. 3. We concluded that the common ancestor of New World and Old World monkeys should have had a smaller and less folded brain than a capuchin, similar to that of a squirrel monkey. The split between small and large New World monkeys occurred about 21 million years ago, from an even smaller common ancestor whose brain should resemble that of an owl monkey: small and barely folded.

**Figure 2.**
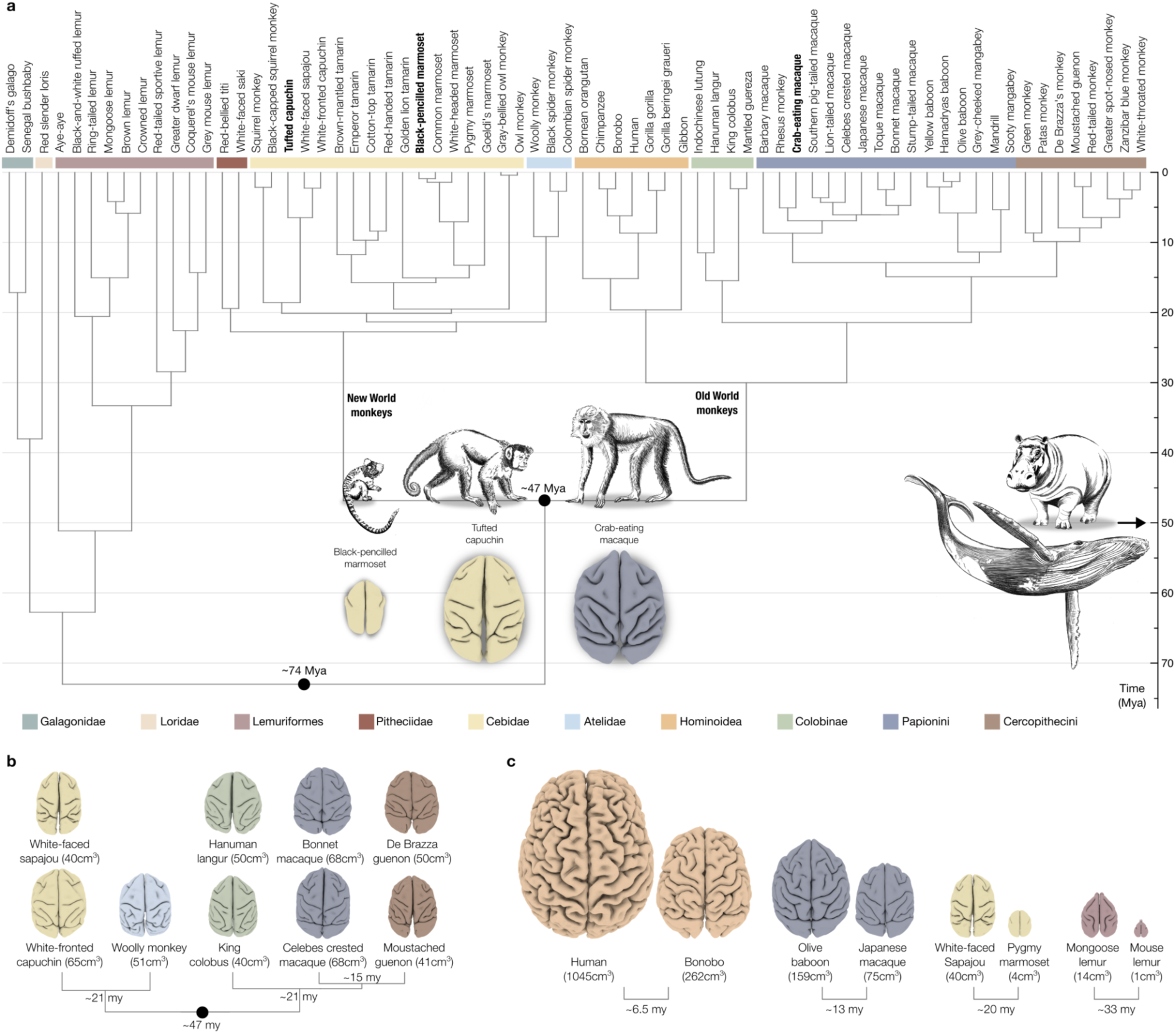
Phylogenetic tree. (a) Phylogenetic tree of the 70 primate species studied, highlighting in particular the branching point of New World and Old World monkeys: macaques and capuchins split from their common ancestor about 47 million years ago – about as long ago as the common ancestor of hippopotamuses and whales lived. Despite this long period of independent evolution, capuchins and macaques evolved strikingly similar brains. The marmoset split from its common ancestor with capuchins much more recently, but evolved a considerably smaller and smoother brain. (b) This is not an isolated example but a recurrent pattern across the phylogenetic tree: brains of similar volume will look strikingly similar even though they evolved independently in different branches, as illustrated here by a sample of brains from different groups including New World and Old World monkeys. (c) To the contrary, brains of different sizes will have very different folding patterns even among species from the same family, as illustrated by a few paired examples. Brain colours indicate different families, as shown in Fig. 1.

**Figure 3.**
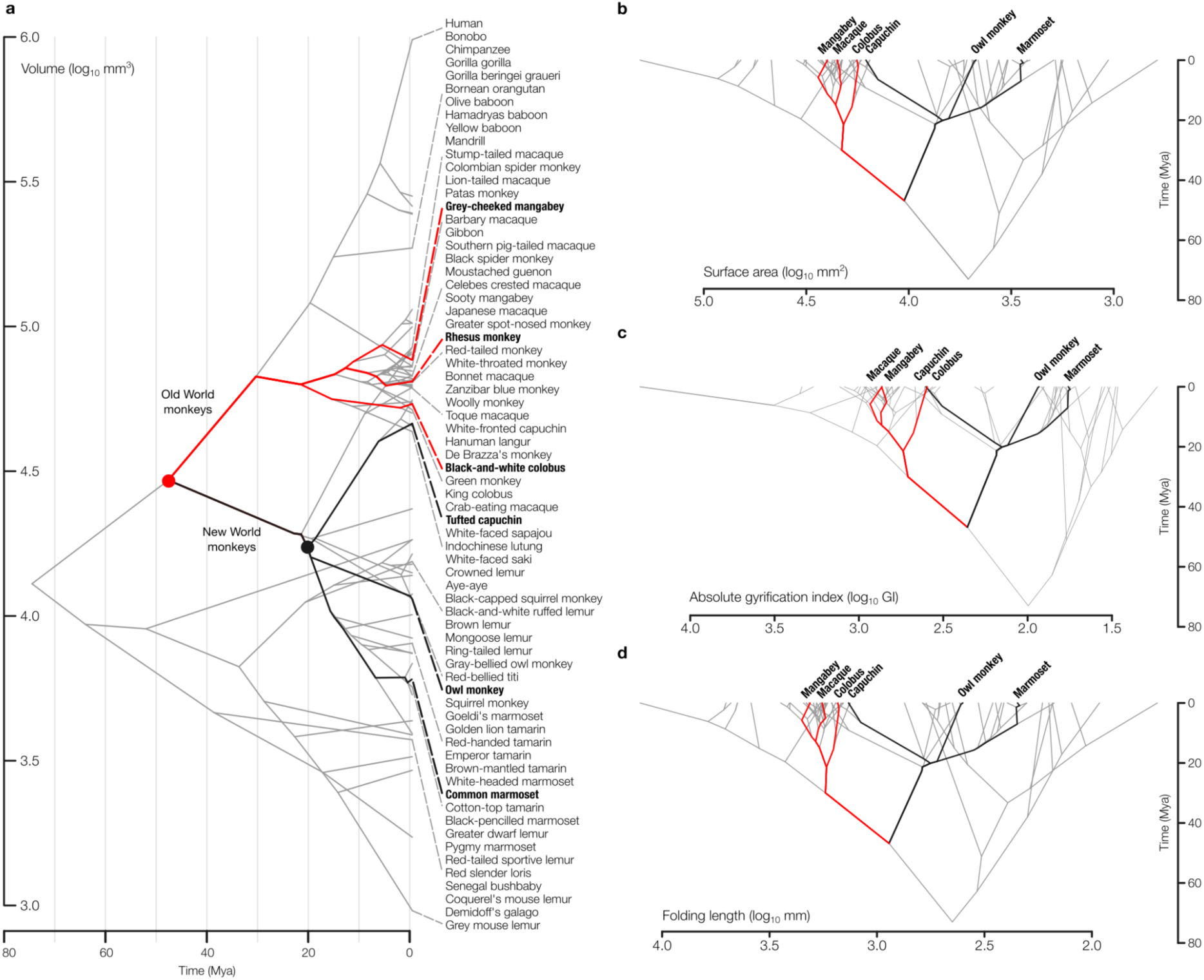
Evolutionary history of brain phenotypes. Phenograms of cerebral volume, cortical surface area and folding evolution across 70 primate species. (a) While in some branches cerebral volume only increases (such as in humans), in others it only decreases (like in grey mouse lemurs). Interestingly, the branch leading to the tufted capuchin shows a mix of increases and decreases. The tufted capuchin has evolved a folding pattern similar to a crab-eating macaque despite the 2 species having split ∼47Mya (red dot), and despite the ancestor of the tufted capuchin being estimated to be small and little folded (∼20Mya, black dot). (b) This pattern is confirmed in the phenogram showing the evolutionary history of surface area (b) where the capuchin independently evolved a surface area very similar to the Old World colobus, macaque and mangabey from a small brained ancestor, together with a very similar amount (c) and length (d) of folding.

Second, we analysed endocranial volume from 179 specimens across 49 New World monkey species and confirmed the result obtained with surface reconstructions from MRI data (Supplementary Fig. S1). Within New World monkeys, the Early-Burst model provided the best fit (an initially high rate of phenotypic change which progressively decreases) (Aristide et al. 2016). Using this model, we estimated the phenotype of the common ancestor of small and large New World monkeys, and confirmed that it had a small brain, similar to that of an owl monkey.

### Primates with similar brain volumes have similar folding patterns, irrespective of phylogenetic similarity

We focused on 6 representative primate species: 3 New World monkeys (marmoset, owl monkey and capuchin) and 3 Old World monkeys (colobus, macaque, and mangabey), illustrated in Fig. 4a. The 3 Old World monkeys had a similar brain volume to the capuchin. Folding patterns are thought to reflect species-specific, genetically encoded patterns of differential gene expression (De Juan Romero et al. 2015). However, a surface morphing shows that the folding pattern of capuchins closely matches that of the Old World monkeys, despite having evolved independently from an almost unfolded common ancestor (Fig. 4b, Video 2). We quantified this similarity by computing curvature maps, and used spin tests to compare the folding maps of our 6 species. Small-brain New World monkeys clustered together (r=0.75, 95% CI=[0.70, 0.80]), and the capuchin clustered with Old World monkeys (Fig. 4c, intra-cluster correlations between r=0.63 and r=0.8, 95% CI=[0.60, 0.81]). The correlation between the folding map of large-brain monkeys (capuchin and Old World monkeys) and small-brain New World monkeys was significantly smaller (from r=0.29 to r=0.5, 95 % CI=[0.23, 0.55]). The difference between the two clusters was statistically significant (z_intra_-z_inter_=0.47, p≪1, Supplementary Fig. S2).

**Figure 4.**
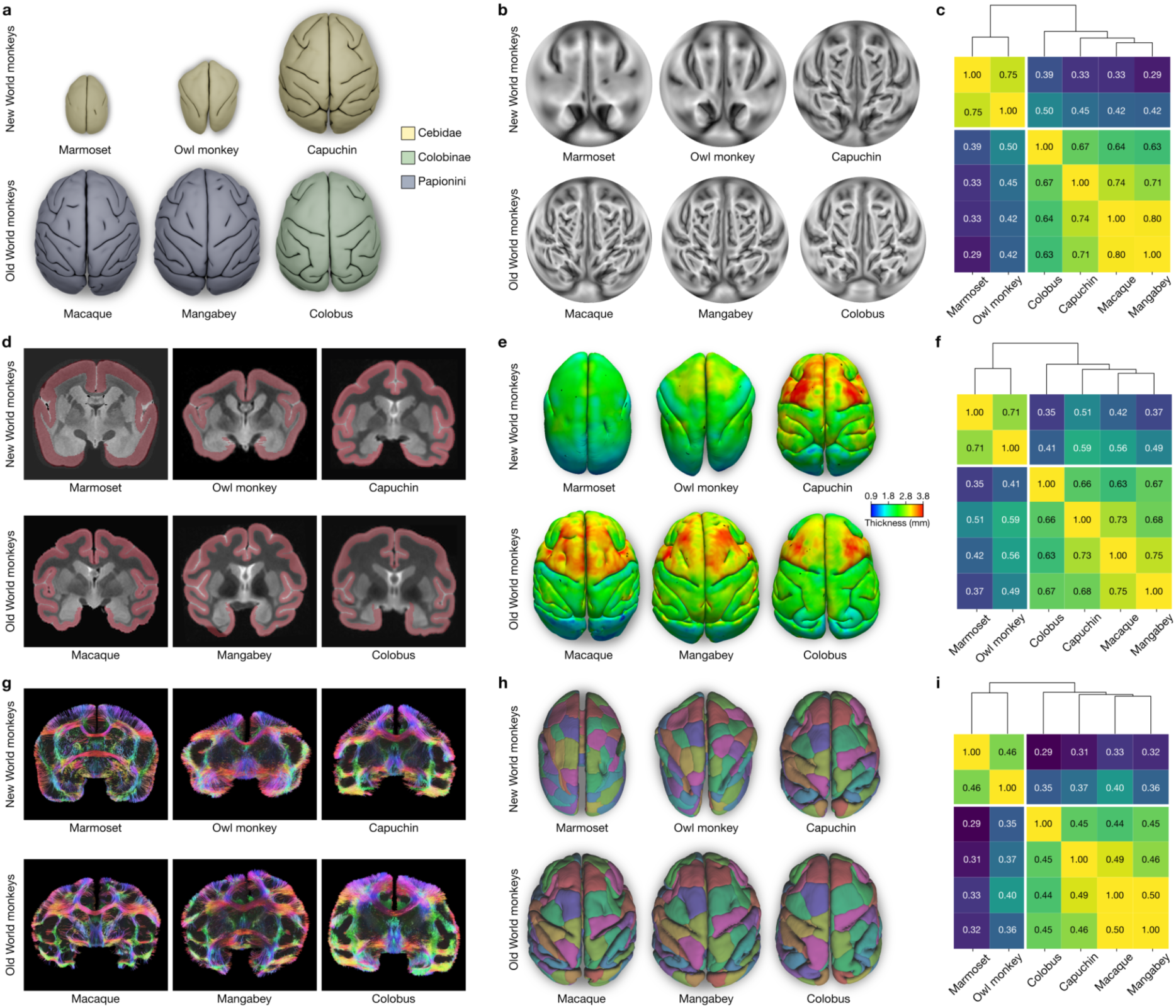
Similarity of folding, thickness, and connectivity patterns across species. Top (a-c): Similarity of folding patterns across species. (a) Dorsal view of 3D brain surface reconstructions of 3 New World and 3 Old World monkeys. Colour indicates phylogenetic relationship, and the surfaces are shown to scale. (b) Flat maps showing the curvature patterns of the cerebral cortices. (c) The correlation matrix shows the similarity of folding patterns across species. Folding patterns were very similar between species with similar brain volume, independent of their phylogenetic relationship. The two species with small brains, marmoset and owl monkey, cluster together with high correlation, as do the 4 species with large brains: capuchin, colobus, macaque and mangabey. Centre (d-f): Similarity of cortical thickness patterns across species. (d) Mid-coronal slices of the 6 brains showing the segmented cortical ribbon in red. (e) Cortical thickness maps shown on the 3D brain surfaces. Brains are not to scale but shown at the same size. (f) The correlation matrix shows the similarity of cortical thickness maps of the different species. Brains with similar volume showed highly correlated thickness maps, independent of their position in the phylogenetic tree. Bottom (g-i): Similarity of connectivity patterns across species. (g) Coronal cut through the reconstructed 3D connections at the level of the anterior commissure. (h) Example of a homologous parcellation across all species. One species’s brain was parcellated into a set number of random regions of equal size (100 parcels used for illustration, 200 were used in the analyses), and this parcellation was then propagated to all other species. (i) The correlation matrix shows the similarity of cortico-cortical connectivity patterns across species. Connectivity patterns are very similar between species with similar brain volumes and similar folding patterns, and less similar between species of the same family but with largely differing brain volume. Hierarchical clustering was applied on the correlation matrices for visualisation.

### Primates with similar brain volumes have similar cortical thickness patterns, irrespective of phylogenetic similarity

We computed cortical thickness maps by segmenting the cortical ribbon in our 6 primate species, illustrated in Fig. 4d. Variations in cortical thickness are an important parameter of neocortical arealisation, and regional differences in cortical thickness are thought to reflect regional differences in proliferation and maturation resulting from patterned gene expression (Lukaszewicz et al. 2005, Reillo et al. 2011, De Juan Romero et al. 2015). However, as it was the case for folding, the thickness map of the capuchin monkey was strikingly more similar to those of phylogenetically distant Old World monkeys with a similar brain volume, than to those of phylogenetically closer New World monkeys with a small brain volume (Fig. 4e). We quantified these differences using spin tests. The cortical thickness maps of small-brain New World monkeys showed a high correlation (r=0.71, 95 % CI=[0.56, 0.82]), whereas the capuchin clustered with Old World monkeys (Fig. 4f, intra-cluster correlations from r=0.63 to r=0.75, 95 % CI=[0.51, 0.83]). The correlations of the capuchin cortical thickness map with its phylogenetically closer relatives were significantly lower (r=0.51 and r=0.59). In general, the correlations between the cortical thickness maps of small-brain and large-brain primates were low (between r=0.35 and r=0.59, 95 % CI=[0.16, 0.71]). The differences between the two clusters were statistically significant (z_intra_-z_inter_=0.34, p≪1, Supplementary Fig. S4 and Supplementary Methods).

### Primates with similar brain volumes have similar connectomes, irrespective of phylogenetic similarity

The different neocortical areas are integrated within a dense network of connections spanning many scales. The precise organisation of these connections is thought to reflect the specific adaptations of each species. In particular, a prominent motif of short associative connections, known as U-fibres, connects neighbouring gyri (Schüz and Braitenberg 2002). If patterned gene expression encodes the pattern of neocortical folds and cortico-cortical connectivity, phylogenetically closer primates should share more similar connectomes than phylogenetically distant ones. However, our analyses of whole-brain tractography from diffusion-weighted MRI showed that once again the connectome of the capuchin monkey matched more closely that of Old World monkeys than those of small New World monkeys (Fig. 4g). To quantify this, we used random parcellations of the neocortex, with varying numbers of parcels, each time starting from one primate and propagating the parcels to the other 5 primates using surface morphing (Fig. 4h). For each random parcellation we built 6 connectomes, one per species (a connectome is a matrix counting the number of connections linking every pair of parcels). We then computed the similarity of the 6 species’s connectomes using a Mantel test. Across thousands of random parcellations, the connectomes of small New World monkeys formed a first cluster of similarity, with the capuchin forming a second cluster together with the Old World monkeys (Fig. 4i). Intra-cluster correlations ranged from r=0.44 to r=0.50, 95% CI=[0.41, 0.52], whereas inter-cluster correlations ranged from r=0.29 to r=0.40. Intra-cluster correlations were significantly higher than inter-cluster correlations (z_intra_-z_inter_=0.15, p≪1, Supplementary Fig. S7).

### Species with similar brain volumes have similar behaviour, irrespective of phylogenetic similarity

Our neuroanatomical findings prompted us to investigate the following question: If we were to predict the behaviour of a species, should we base our prediction on the behaviour of species with a similar neuroanatomy, or on the behaviour of species phylogenetically close? We collected 20 behavioural phenotypes from the literature for our 70 primate species. For each species we built a behavioural vector (39 dimensions, including phenotypes such as tool use, social learning and locomotion), and a neuroanatomical vector (5 dimensions, including measurements such as total brain volume, surface area and gyrification index). We used these vectors to compute behavioural and neuroanatomical similarity matrices, along with a phylogenetic similarity matrix obtained from the phylogenetic tree (in the Brownian Motion model, the phylogenetic distance between two species is the length of the phylogenetic tree branches from one species to the other). Our results showed a strong correlation between neuroanatomical similarity and behavioural similarity (r=0.56, p<0.001, 95% CI=[0.49, 0.63], Fig. 5a), whereas the correlation between phylogenetic similarity and behavioural similarity was significantly lower (r=0.29, p<0.001, 95% CI=[0.18, 0.40], Fig. 5b). We used partial correlations to further test for an effect of neuroanatomy independent of phylogeny, and found a significant correlation of neuroanatomical and behavioural similarity when controlling for phylogenetic similarity (r=0.50, p<0.001, 95% CI=[0.42, 0.58]). By contrast, the correlation between phylogenetic and behavioural similarity when covarying neuroanatomical similarity was not significant (r=0.03, p=0.63, 95% CI=[-0.10, 0.16], Fig. 5d). Hence, neuroanatomical similarity appeared as a significantly stronger predictor of behavioural similarity than phylogenetic similarity.

**Figure 5.**
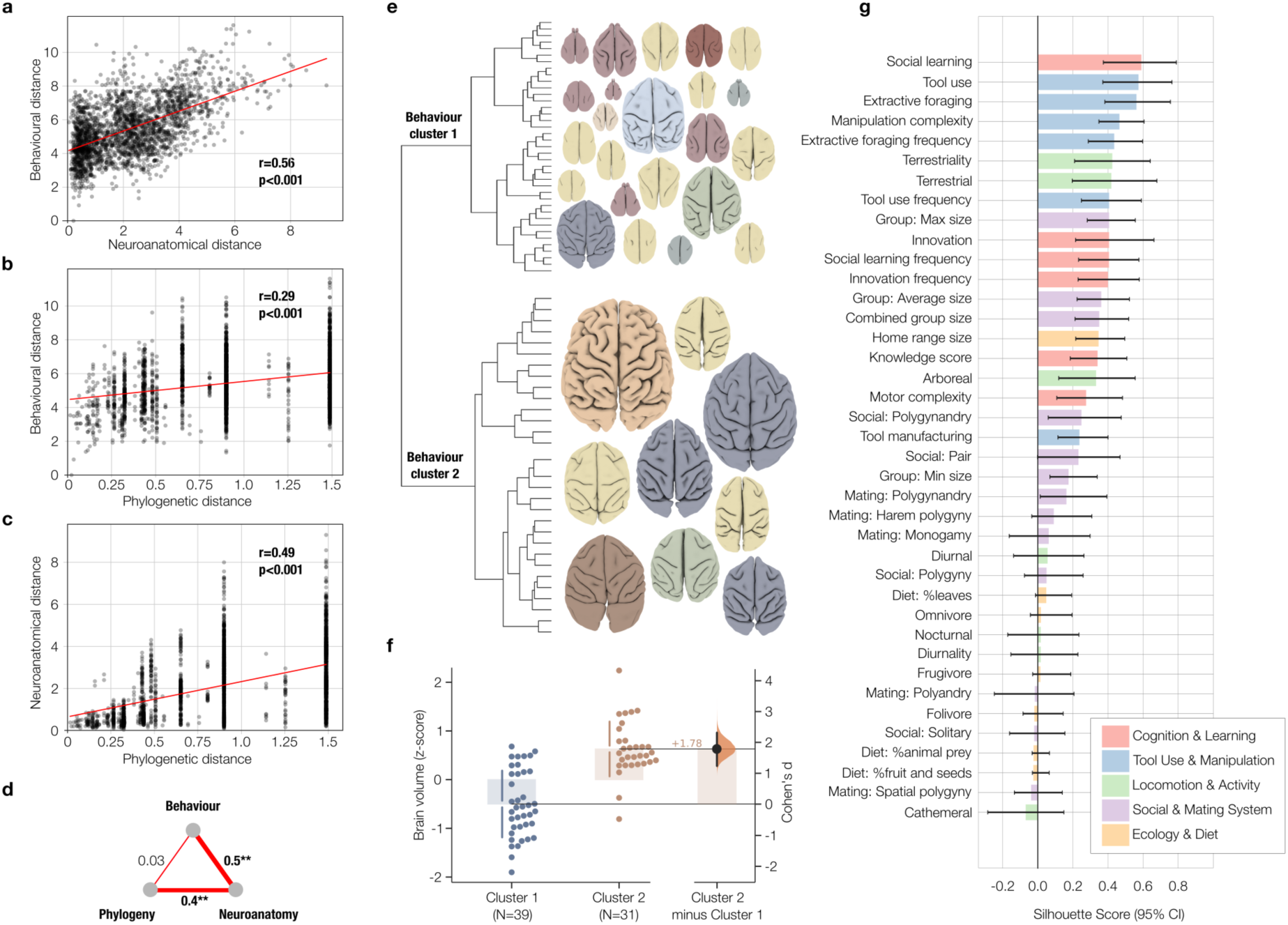
Similarity of behaviour across species. The correlation between neuroanatomical similarity and behavioural similarity was strong (panel a, r=0.56, p<0.001, 95% CI=[0.49, 0.63]) and significantly larger than the correlation between phylogenetic similarity and behavioural similarity which was much lower (panel b, r=0.29, p<0.001, 95% CI=[0.18, 0.40]. Neuroanatomical and phylogenetic distance were also correlated (panel c, r=0.49, p<0.001, 95% CI=[0.40, 0.56]), but partial correlations a significant correlation of neuroanatomical and behavioural similarity when controlling for phylogenetic similarity (panel d, r=0.50, p<0.001, 95% CI=[0.42, 0.58]) and no significant partial correlation between phylogenetic and behavioural similarity when controlling for neuroanatomical similarity (panel d, r=0.03, p=0.63, 95% CI=[-0.10, 0.16]). (e) Hierarchical clustering based only on behaviour splits the dataset into two clusters with significantly different cerebral volume). Brain surface examples for the 2 clusters are shown to scale and coloured by phylogenetic relationship. (f) Significant difference in brain volume between the behavioural clusters 1 and 2 (Cohen’s d: 1.78, p≪1). (g) Importance of the different behavioural variables in the clustering (silhouette scores), with the categories cognition and learning and tool use and manipulation ranking highest (error bars show 95% confidence intervals).

Hierarchical clustering based on behaviour confirmed this result: The first level of the hierarchy clustered species from different branches of the phylogenetic tree and reflected well their neuroanatomical similarity (Fig. 5e) showing a significant difference in brain volume between the two behavioural clusters (Cohen’s d: 1.78, p≪1, Fig. 5f). The importance of different behavioural phenotypes for the clustering was assessed using silhouette scores (Fig. 5g). The behaviours most strongly different between the 2 main clusters were social learning, tool use, extractive foraging, manipulation complexity, terrestriality, innovation and group size (mean silhouette coefficient between 0.35 and 0.59). The least different behaviours included diet composition and activity period (mean silhouette coefficient between 0.06 and −0.07). Neuroanatomical phenotypes were all very different between the two clusters (mean silhouette coefficient between 0.32 and 0.35).

## Discussion

We describe a striking case of convergent evolution in neuroanatomical organisation and behaviour, appearing independently across distant branches of the primate phylogenetic tree. These similarities are not readily explained by a framework where patterns of neocortical folding, thickness, and cortico-cortical connectivity are shaped solely by genetically encoded patterns and experience-dependent processes.

We will first consider several alternative hypotheses and then propose that the integration of mechanical morphogenesis – the emergence of complex shapes from mechanical instabilities (Foubet et al. 2019, Heuer and Toro, 2019) – provides a more parsimonious explanation for our findings. Finally, we will discuss the broader implications of this new integrated framework for our understanding of the development and evolution of brain organisation.

A first possibility could be that the convergent evolution we observed results from similar selective pressures. However, the existence of a strong pressure on precise folding patterns would be very surprising. The brain’s remarkable plasticity allows it to adapt even to substantial neuroanatomical changes – such as agenesis of the corpus callosum or even the dysgenesis of an entire brain hemisphere – which can sometimes be compensated, producing only mild or barely detectable symptoms (Paul et al. 2007, Muckli et al. 2009). Given this plasticity, the convergent evolution of a genetic encoding for folding patterns, cortical thickness, and cortico-cortical connectivity seems highly unlikely.

Second, it could be that the phylogenetic tree distances we used are an unreliable proxy for genetic similarity. However, the primate phylogenetic tree is notably stable, and different approaches to its construction lead to very similar results in the topological relationship between species as well as divergence times (Arnold et al. 2010, Dos Reis et al. 2018, Vanderpool et al. 2020, Lemoine and Gascuel 2021, Kumar et al. 2022). Additionally, the study of the phylogenetic tree of placental mammals shows that tree structure is consistent no matter if only coding or non-coding regions are used for its construction (Foley et al. 2023). The tree we used was built based on genome alignment (Dos Reis et al. 2018), however the results were identical with a tree built from 11 mitochondrial and 6 autosomal genes (Arnold et al. 2010, Supplementary Fig. S11).

Third, our conclusion depends on the pattern of neocortical folding, thickness and connectivity in capuchins having evolved from an ancestor with a small, largely unfolded brain. We obtained this ancestral estimation by fitting phylogenetic evolutionary models to MRI and endocast data from 105 primate species. Our conclusion is also supported by the available fossil record. The endocast of *Chilecebus carrascoensis* is thought to be a good representation of the common ancestor shortly after the divergence of New World and Old World monkeys (Ni et al. 2019). Although endocasts only provide an imprint of the outer surface of the brain but not a precise reconstruction of folding patterns, the correlation between brain volume and degree of folding in primates is very strong (r=0.95, CI=[0.92, 0.97] in our 70 primate species, see also Rogers et al. 2011, Heuer et al. 2019). With an estimated brain volume of about 8 cm^3^, *Chilecebus carrascoensis* should have been similar to today’s marmosets or tamarins: small and significantly less folded than capuchins (Powell and Barton 2017, Heuer et al. 2019, Ni et al. 2019). For the common ancestor to have a degree of folding similar to that of capuchins and macaques, its brain would have to be almost 10 times larger than that of *Chilecebus carrascoensis*.

Fourth, we could also consider the possibility that the common ancestor of New World and Old World monkeys had the genetic encoding of the folding and connectivity patterns of capuchins and macaques, but in a hypothetical “dormant” state. Possibly, this dormant pattern would only become apparent given enough brain growth. However, the notion that folding patterns would not emerge, but be inherited from a common ancestor imposes significant theoretical constraints. Numerous Old World monkeys have larger and more folded brains than macaques, for example, humans and other great apes. Would the dormant ancestral organisation contain the full possibilities of future neuroanatomies up to humans? What type of selective pressure would make this preemptive implementation of such a complex phenotype possible? Should we suppose that capuchins and macaques inherit an ancestral organisation, but a different mechanism leads to the continued elaboration of brain organisation in great apes and humans?

Overall, our results highlight a limitation of the hypothesis that neocortical folding, thickness and connectivity patterns result from a genetically encoded pattern of differential expression. This hypothesis comes from experimental results involving a variety of genetic perturbations. Genetic mutations, over- and underexpression of genes involved in neurogenesis, migration, or axonal guidance, can lead to changes in folding (reviewed by Fernández and Borrell 2023, Akula et al. 2023). But how is this initial pattern encoded? A potential mechanism could involve signalling molecules such as Fgfs, Wnts and Shh, as suggested by O’Leary et al. (2007, 2013), and similar to the “French Flag” model of Wolpert (1969): Gradients of such signalling molecules would be produced from a reduced number of centres; each position on the cortex would be characterised by a unique concentration; a decoding mechanism would then control the differential expression of specific sets of genes, leading to the formation of a species specific pattern of neocortical areas, as well as a pattern of gyri and sulci. Similarly, the decoding mechanism could regulate the expression of chemorepulsive and chemoattractive molecules, leading to the formation of specific cortico-cortical connections. For a small brain such as that of a mouse or a small lemur, with few neocortical areas and mostly transcallosal cortico-cortical connections, the encoding of areas, folding and connectivity patterns could be relatively simple. But a complete mouse brain fits within a single human neocortical fold: As brain size increases, the number of neocortical areas increases linearly and the number of connections exponentially – the genetic encoding of folding, thickness and connectivity patterns would become increasingly complex, and the likelihood of a classical convergent evolution would rapidly decrease.

Instead, we identified a fundamentally different kind of convergence – one driven by mechanical constraints rather than genetic adaptation. We propose that a 3-factor framework, integrating mechanical morphogenesis together with genetics and experience-dependent processes, offers a more parsimonious account of our findings. In essence, the mechanical morphogenetic process that leads to folding pattern formation rests on two rather uncontroversial assumptions: (1) the development of the neocortical neuropil after the end of neuronal migration leads to the rapid expansion of the neocortex without a similar expansion of the white matter core (Rash et al. 2022), (2) cortex and core have similar stiffness – softer than jelly – and are relatively incompressible (Budday et al. 2015). Under these conditions, neocortical expansion will initially be accommodated by neocortical compression and extension of the core. If growth continues, a critical point will be reached where a neocortical folding pattern emerges, allowing for neocortical expansion without further expansion of the core. In addition to neocortical expansion, the critical point depends importantly on cortical thickness: a thin cortex, like in the cerebellum, will fold earlier and with smaller folds than a thicker one. If expansion is insufficient or if the cortex is too thick, there will be no folding. The pattern of these folds will be determined by its intrinsic wavelength, and regulated by the initial shape of the brain. This basic process has been extensively described and simulated (Todd 1982, Toro and Burnod 2005, Toro 2012, Bayly et al. 2013, Tallinen et al. 2014, Tallinen et al. 2016, see Kroenke and Bayly 2018 for a review).

Comparative studies of the cerebral and cerebellar cortex provide further information to understand the 3-factor framework. They reveal a stark dissociation between two groups of neuroanatomical traits: First, a group of “stable” traits, such as cortical thickness and folding wavelength, that show little variation from mice to whales. Second, a group of “diverse” traits, such as total brain volume or the tangential extension of the neocortex, which vary over 4 orders of magnitude (Heuer et al. 2019, Heuer et al. 2023). This split hints at two different genetic architectures. First, conserved pathways involving major genes responsible for the fundamental processes leading to the formation of a mammalian cortex (including symmetric and asymmetric progenitor division, proliferation of radial glia, derivation into neurons, migration). During normal development, these processes lead to a cortex of a stable thickness and similar mechanical properties across species (Budday et al. 2019, Iwashita et al. 2020). These processes seem to be the main target of the current genetic manipulations (reviewed by Llinares-Benadero and Borrell 2019, Fernández and Borrell 2023, Akula et al. 2023). Second, a strongly polygenic architecture, aggregating thousands of infinitesimal effects in regulatory regions supporting the variability of the diverse phenotypes (Toro et al. 2014, Biton et al. 2020, Grasby et al. 2020, van der Meer and Kaufmann 2022). The paradigmatic example of such a phenotype is human height – a quantitative trait influenced by >20% of the genome (Yengo et al. 2022). These polygenic effects seem to be shared among diverse phenotypes, leading to strong allometric patterns (Finlay et al. 2001, Heuer et al. 2023).

In summary, we explain the neuroanatomical and behavioural similarity between capuchins and macaques as follows: (1) a conserved genetic program would lead to the formation of a cortex with a stable thickness and mechanical properties, (2) strongly polygenic effects would lead to a comparable degree of neocortical expansion, (3) this expansion would be sufficient to make folding emerge (a mechanical morphogenetic effect), (4) the large-scale geometric and mechanical constraints introduced by folding would result in a feedback on cellular maturation, cortico-cortical connectivity and behaviour. Finally, (5) experience would further sculpt this anatomy throughout life. Conversely, the reduced folding of marmosets and tamarins would be attributed to insufficient neocortical expansion (a polygenic effect). Our study is, however, not about a single case of convergence between a few species, but reveals a fundamental organising principle of brain morphology (as shown in Video 1). In general, the strongly conserved processes shared by all primates – for example, a neocortical structure of similar thickness and mechanical properties – combined with the highly diverse degree of cortical expansion would explain the emergence of similar folding patterns across primates. Furthermore, this should be observed at the level of species but also individuals: the folding pattern of a small capuchin, for example, should approach that of a large squirrel monkey (note that in humans, large brains are disproportionately more folded than small ones, Toro et al. 2008). Combined, the stable radial structure of the cortex, the polygenic increase in tangential expansion and emergent mechanical morphogenetic processes, provide a parsimonious explanation for the convergence of neuroanatomical phenotypes of primates across the phylogenetic tree.

In conclusion, progress in understanding brain development and evolution should come from fully integrating mechanical morphogenesis alongside genetics and experience-dependent processes into a 3-factor framework. A growing body of research suggests that mechanical forces can impact key biological processes, such as cell proliferation, apoptosis, cell fate, gene expression and axonal guidance (Saha et al. 2008, Le et al. 2016, Kumar et al. 2017, Thompson et al. 2019, Oliveri et al. 2021, Wagh et al. 2021, Schaeffer et al. 2022). This type of feedback could be at the root of the convergence in cortical thickness, connectivity patterns and behaviour. It has been shown, for example, that the cytoarchitecture and electrophysiology of motor and somatosensory areas in capuchin monkeys resemble more closely those of macaques than those of smaller New World monkeys (Padberg et al. 2007, Mayer et al. 2019). Mechanical morphogenesis provides a way through which “more” becomes “different” (to paraphrase W. A. Anderson,1972). This comprehensive approach not only provides a more parsimonious explanation for our findings, but also opens an exciting new perspective for the study of brain development and evolution.

## Acknowledgements

KH and RT were funded by project NeuroWebLab (ANR-19-DATA-0025), DMOBE (ANR-21-CE45-0016), and the European Union’s Horizon 2020 research and innovation programme under the Marie Skłodowska-Curie grant agreement No101033485 (KH Individual Fellowship). For the purpose of open access, the author has applied a CC-BY public copyright licence to any Author Manuscript version arising from this submission. This work is supported by the ERC grant (UNFOLD, ERC-2023-SyG n°101118729). Funded by the European Union. Views and opinions expressed are however those of the author(s) only and do not necessarily reflect those of the European Union or the European Research Council Executive Agency. Neither the European Union nor the granting authority can be held responsible for them. The MNHN gives access to the collections in the framework of the RECOLNAT national Research Infrastructure. RBM was supported by the Biotechnology and Biological Sciences Research Council (BBSRC) UK [BB/N019814/1], the Wellcome Centre for Integrative Neuroimaging is supported by core funding from the Wellcome Trust [203139/Z/16/Z]. SN was supported by Fonds de recherche du Québec – Nature et technologies (FRQNT), number: 353575.

## Author contributions

KH and RT conceptualised the study, administered and supervised the project, acquired funding, created the visuals, and wrote the original draft of the manuscript. KH, RT, NT and LA designed the software and methodology, performed the analyses. KH, RT, SFA, SN, RM collected and curated the data. LA, MH, RBM, TS, MS contributed brain tissue, CT and MRI data. All authors contributed in reviewing and editing the final manuscript.

## Competing interests

The authors declare no competing interests related to the present work.

## Methods

### 1. Data

#### Structural MRI

We collected structural brain MRI data from 70 primate species, comprising 108 specimens, from sources including: the National Chimpanzee Brain Resource (Rilling and Insel 1999), the Brain Catalogue Primates project (Heuer et al. 2019), the Japan Monkey Center (Sakai et al. 2018), Primate Brain Bank (through Navarrete et al. 2018), the Primate Data Exchange (Prime-DE, Milham et al. 2018, 2020, 2022) and ABIDE (Di Martino et al. 2013). MRI data for 5 new species from the wet comparative anatomy collection of the National Natural History Museum (MNHN) were scanned at the Center for Neuroimaging Research (CENIR) of the Institut du Cerveau et de la Moelle Épinière (ICM, Paris, France). High resolution MRI images were acquired using either a 3T Siemens Tim Trio system, a 3T Siemens Prisma, or an 11.7T Bruker Biospec. Each dataset was acquired with a 3D gradient-echo sequence (FLASH). Parameters (Field of View, Matrix size, TR, TE) were adjusted so as to obtain the highest resolution possible with our scanner (from 100 to 450 μm isotropic). TR and TE were always chosen as minimum. Flip angle was fixed to 20° at 3T and 10° at 11.7T. The number of averages was chosen to maintain a scanning time below 12 h. The data encompasses all major families of primates, except for one pro-simian (non-monkey) family, tarsiers, and provides a great sample across the primate phylogenetic tree from the smallest to the largest primate brain, spanning an 800-fold difference in brain volume (see Supplementary Table 4 for a detailed list).

#### Brain endocasts

We collected brain endocast data from 179 specimens across 49 New World monkey species (Aristide et al. 2016). The skulls were digitised using X-ray computed tomography or micro-computed tomography. 3D virtual endocasts were created using a threshold-based 2D segmentation procedure. The data covered all major 17 genera and include about half of the extant platyrrhines and represent a large sample of the platyrrhine phylogeny (see Supplementary Table 5 for a detailed list).

#### Diffusion-weighted MRI

We collected high resolution diffusion weighted MRI data for 6 primate species: 3 New World monkeys (common marmoset, grey-bellied owl monkey, tufted capuchin) and 3 Old World monkeys (black-and-white-colobus, rhesus macaque, grey-cheeked mangabey) from sources including: the Digital Brain Bank through Bryant et al. (2021) and Folloni et al. (2019), respectively, and the Marmoset Brain Mapping project (Liu et al. 2021). The datasets were acquired at 7T field strength, using multi-shell diffusion schemes with b-values of 2400, 4800, 7200 s/mm2 (marmoset), and 4000 s/mm2 (all other primates), and 126 and 128 directions for the marmoset and the other primates, respectively. Images were acquired *ex-vivo* with isotropic resolutions of 0.15 mm in the marmoset, 0.4 mm in the owl monkey, 0.5 mm in the macaque, and 0.6 mm in the cebus, colobus, and mangabey.

#### Behavioural data

We collected behavioural data from the scientific literature and large online databases, including AnAge (https://genomics.senescence.info/species/index.html), Encyclopedia of Life (https://eol.org, Parr et al. 2014), Animal Diversity Web (https://animaldiversity.org, Yahnke et al. 2013), PanTHERIA (Jones et al. 2009), and NCBI Taxonomy (https://www.ncbi.nlm.nih.gov/taxonomy). Our list contained 20 behavioural variables, such as group size, activity period, whether they use or manufacture tools and their reported manipulation complexity, innovation, social learning, food resource and diet composition, mating and social systems, habitat type (whether they are arboreal or terrestrial), home range size, etc. (Bentley-Condit and Smith 2010, Jaeggi and Van Schaik 2011, Reader et al. 2011, Dubman et al. 2012, Heldstab et al. 2016, Schuppli et al. 2016, Decasien et al. 2017, Powell and Barton 2017). An overview of all collected variables can be found in Supplementary Table 3.

#### Endocranial and brain volume data

We also collected data on endocranial volume and brain volume from the literature (Dubman et al. 2012, Decasien et al. 2017, Powell and Barton 2017).

### 2. Surface reconstruction and neuroanatomical measurements

We used several tools to produce cortical surface reconstructions from the MRI data. We reoriented the data to align the brains with respect to the stereotaxic axes using Reorient (https://neuroanatomy.github.io/reorient, Heuer et al. 2020). A first mask was obtained interactively using Thresholdmann (https://neuroanatomy.github.io/thresholdmann, Heuer et al. 2024), which allows the creation of space-varying segmentation thresholds. These masks were then refined using BrainBox (https://brainbox.pasteur.fr, Heuer et al. 2016). BrainBox is a Web application providing several tools for collaborative manual segmentation of MRI data. The main task in BrainBox was to make sure that sulcal regions were open, and to correct additional segmentation errors. The masks generated by BrainBox were then used to produce surface meshes, using the implementation of the marching cubes algorithm provided by Scikit-Image (van der Walt, 2014). The meshes were then regularised using Graphite (https://github.com/BrunoLevy/GraphiteThree), which also automatically corrects a certain number of topological errors. Finally, the topology of the meshes was manually edited to obtain meshes of genus 0 (topologically spherical) using MeshSurgery (https://github.com/neuroanatomy/MeshSurgery).

A series of neuroanatomical measurements were obtained from these meshes, including brain volume, cerebral volume, surface area, and gyrification index, using the command line tool meshgeometry (https://github.com/neuroanatomy/meshgeometry).

### 3. Phylogenetic comparative analysis of MRI data and ancestral trait estimation

We obtained phylogenetic tree data from Dos Reis et al. (2018), which provides divergence times and topological relationships for the primate phylogeny based on phylogenomic data. We used the R package mvMORPH (Clavel et al. 2015) to fit 3 different evolutionary models: (1) the Brownian Motion model, where phenotypes vary randomly through time, (2) the Ornstein-Uhlenbeck model, where in addition to random variation phenotypes tend toward an advantageous value, and (3) and Early-Burst model, where the rate of change slows down from the beginning to the end of the evolutionary process. We selected the best fitting model using the Akaike Information Criterion corrected for small sample size (Supplementary Table 1). The best fitting model was the Brownian Motion model. Ancestral phenotypes were hence estimated using the Brownian motion model. The estimated ancestral phenotypes were then used to identify the extant species whose brain measures provided the closest match to the estimated phenotypes of the common ancestor of New World and Old World monkeys, and Cebidae with large and small brains. For this, we computed the Euclidean distance between the vector of estimated ancestral phenotypes and the phenotypes of all species in our sample.

### 4. Endocasts reconstruction and measurements

A large sample of virtual cranial endocasts from 49 New World monkey species was used to estimate ancestral phenotypes in an additional independent dataset. 3D reconstructions of the endocasts of 179 specimens across 49 species were obtained from Aristide et al. (2016, Supplementary Fig. S1). We obtained measures of endocranial volume, the volume enclosed by the cranial endocast surface, for all 179 endocasts.

### 5. Phylogenetic comparative analysis of endocast data

The phylogenetic tree was obtained from Aristide et al. (2016) and is based on the fossil-calibrated Bayesian molecular tree presented in Aristide et al. (2015). We used ape (Paradis 2012), phytools (Revell 2011), Rphylopars (Goolsby et al. 2016), pulsR (https://github.com/Schraiber/pulsR) and mvMORPH (Clavel et al. 2015) to perform phylogenetic analyses, test the fit of different evolutionary models and estimate ancestral phenotypes. We tested the Brownian Motion model, the stable model, which is similar to the Brownian Motion model but allows for different rates of variation along different branches, the Ornstein Uhlenbeck model, and the Early Burst model. As we include several specimens per species, the mean standard error was used in the model fit. Based on the Akaike Information Criterion values, we selected the model that provided the best fit for the platyrrhine data sample (Supplementary Table 2).

### 6. Surface morphing

For a quantitative comparison across the phylogeny we developed a method to establish geometric homologies across species. We used a non-linear surface matching algorithm which builds a mapping from one native surface to the other using a reduced number of eigenvectors from the source mesh Laplacian. A linear combination of these eigenvectors was found which minimised metric distortions while maintaining smoothness (see Supplementary Methods I.2 for more information). We used this surface morphing method to quantitatively compare patterns of cortical folding, thickness, and connectivity across species.

### 7. Random surface parcellations

We computed random parcellations of the cortical surface meshes into a fixed number of parcels. The centroids of the parcels were obtained from a Poisson disk sample set of vertices generated with a sample elimination method (Yuksel 2015), so that parcels were of comparable size.

### 8. Cross-species comparison of the folding pattern

Cortical mean curvature was computed for all brains using meshgeometry (https://github.com/neuroanatomy/meshgeometry). Using the aligned surfaces, we computed pairwise correlations using Pearson correlation between curvature maps of 2 species, and the spin test method to assess their significance (Vos de Wael et al. 2020). See Supplemental Methods I.3 for more information.

### 9. Cross-species comparison of cortical thickness

#### Computation of cortical thickness

We automatically segmented the cortical ribbon across all species by detecting the points of maximum change in grey level (see Supplementary Methods I.4) and derived cortical thickness maps. The automatic cortical segmentations were visually inspected and manually refined using BrainBox (Heuer et al. 2016). This refinement was projected back to the cortical thickness map. To assess the validity of our thickness estimation, we compared the thickness map obtained for our rhesus macaque to the Yerkes macaque thickness map (Hayashi et al. 2021, Supplementary Fig. S3). The two methods provided very similar results (r=0.74, p≪1).

#### Comparison of cortical thickness maps

For each species, we projected the brain surface mesh to a sphere (Toro and Burnod 2003) and resampled the thickness map to the subdivided icosahedron (5 subdivisions) of the sphere of each of the two brains. We then computed Pearson correlation between the thickness maps across species. Finally, we used the spin test method (1000 spins), as well as the sampling distribution and confidence intervals to assess the significance of the correlations (see Supplementary Methods I.4 and Supplementary Fig. S4). We applied hierarchical clustering on the correlation matrix for visualisation of the correlations.

### 10. Cross-species comparison of connectomes

#### Whole-brain tractography

The diffusion-weighted imaging data were preprocessed using FSL (Jenkinson et al. 2012) including correction for eddy current, motion, and susceptibility artefacts (see Ardesch et al. 2021 for details). We performed whole-brain deterministic tractography using Mrtrix (Tournier et al. 2019) (maximum fibre length adapted for each species, termination of tracking below tensor FA value 0.02, step size adjusted to data resolution). To capture connectivity throughout the white matter as well as into the cortex, we tracked with a low tensor FA value to terminate the fibre tracking (Guevara et al. 2020) and used the cerebral masks for tracking.

#### Computation of connectomes

We computed whole brain connectomes for the 6 primate species using parcellations with 200 parcels. To exclude an effect of parcellation size on our results, we repeated our analyses using parcellations with 50, 100, 150, 200, 250 and 300 parcels. Our conclusions were not affected by the number of parcels (Supplementary Methods I.5). To exclude an effect of the species used to create parcellations, we computed these parcellations in each brain, and then propagated them to all other brains. This enabled us to have homologous regions across species and compare patterns of brain organisation based on these matched parcels. For robustness, we generated a set of 100 random cortical surface parcellations per species and per number of parcels, and propagated these parcels to all other species. In this way, we obtained 6 sets of 600 matching parcellations across all species for the different numbers of parcels (Supplementary Fig. S5). Based on the 6 sets of 600 homologous parcellations for all species generated using our method described in section 7, we computed 600 connectomes per species using Mrtrix (Tournier et al. 2019).

#### Connectome comparisons

We used Spearman correlation to compare the corresponding connectomes of 2 species for each of the 600 parcellations, and then computed the mean correlation matrix between pairs of species. We used a Mantel test (100 iterations for each of the 600 parcellations) to estimate the significance of the correlations (see Supplementary Fig. S7). We applied hierarchical clustering on the mean correlation matrix for visualisation of the correlations.

### 11. Cross-species comparison of behavioural variables

Behavioural variables were log-converted if that made them more normally distributed (goodness-of-fit test), and were then mean-centred and standardised. Missing values were imputed using an iterative imputation algorithm (Pedregosa et al. 2011, Abraham et al. 2014). For each species, we built a behavioural vector (39 dimensions, including tool use, social learning, locomotion, etc.), and a neuroanatomical vector (5 dimensions, including total brain volume, surface area, gyrification index, etc.). We used these vectors to compute pairwise correlations and converted these into distances to obtain behavioural and neuroanatomical distance matrices. Additionally, we converted the phylogenetic tree into a distance matrix (in the Brownian Motion model, the phylogenetic distance between two species is proportional to the length of the phylogenetic tree branches from one species to the other). We then used Mantel tests (10.000 permutations) to determine whether the neuroanatomical, behavioural and genetic distances were correlated.

## Data availability

All data analysed in this study is openly available from the sources specified in the manuscript.

## Code availability

Code for reproducing our statistical results and our figures will be openly available on GitHub once the manuscript has been accepted.

## I. Supplementary Methods

### I.1 Ancestral endocranial volume estimations

**Figure S1.**
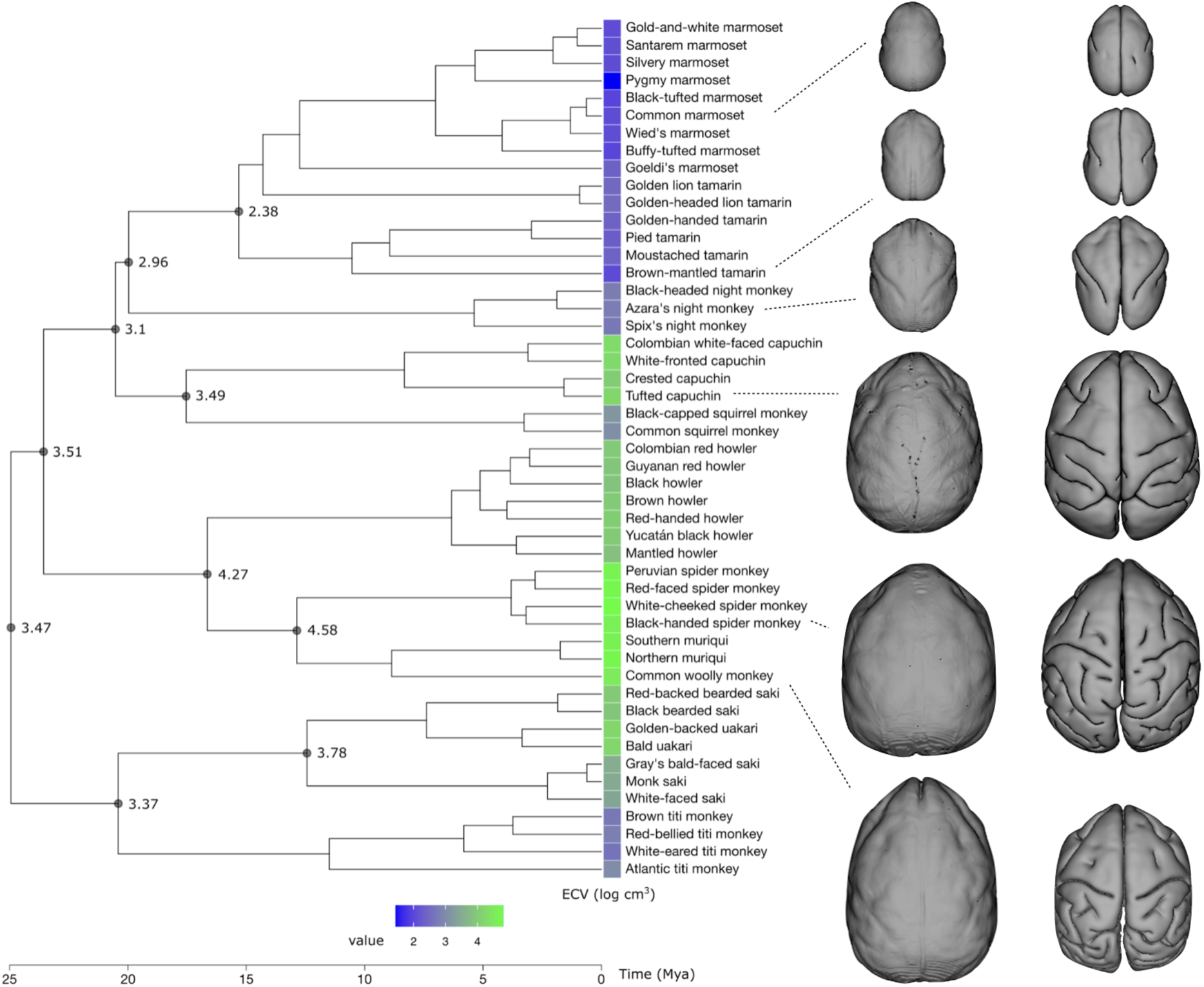
Ancestral endocranial volume estimations. Left: Phylogenetic tree of 49 New World monkeys with ancestral endocranial volume estimates. Estimates are based on the Early Burst model which had provided the best fit to the data when testing different evolutionary models. Using endocasts from 179 specimens across 49 platyrrhine species, the common ancestor of New World monkeys has been estimated to have had a small ECV, and the ECV of the ancestor of Cebidae with large and small brains has been estimated to be even lower, comparable to an owl monkey, who have rather small and little folded brains. Right: 6 examples of endocasts from small and large-brained species, including the marmoset, owl monkey and capuchin, show the imprint of the folding pattern and confirm the relationship between volume and folding.

### I.2 Surface morphing

For a quantitative comparison across the phylogeny we developed a method to establish geometric homologies across species based on nonlinear surface matching. We obtained a trajectory for neocortical expansion across primate species going from small lemurs towards great apes (Video 1) bringing all primate species into a continuous space. This allowed us to quantitatively compare patterns of cortical folding, thickness, and connectivity across species, and to obtain sets of 600 matching surface parcellations across all species (Figure S5, Videos 1 and 2).

For building a homology between two meshes *M*_1_ and *M*_2_ we first compute Laplacian eigenvectors of the source mesh *M*_1_ solving *Lv* = λ*v*, where *L* is the cotangent Laplacian, *v* the eigenvectors, and λ the eigenvalues. A deformation field *D* is then defined as the linear combination of *k* eigenvectors with the smallest eigenvalues,

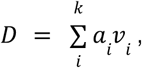

which produce an orthogonal basis for expressing a smoothly varying function over the mesh (typically *k* = 100). Each vertex *p* in *M_1_* is projected into a point *m* in *M_2_* using spherical deformations of each mesh.

The projection function *f* includes also the deformation field *D*, such that

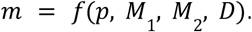

A deformation cost function is defined which compares the length of each edge in the native mesh *M*_1_ with that of its projection in the native mesh *M*_2_:

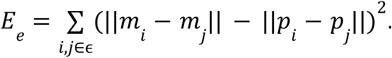

Additionally, we include an energy term related to the smoothness of the deformation:

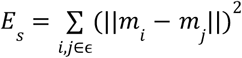

This cost function is minimised using the L-BFGS algorithm to solve for *a*: the coefficients of the linear combination of eigenvectors defining the deformation field *D*, where the weightings were α = 10 and β = 1:

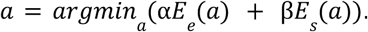

### I.3 Cross-species comparison of the folding pattern

#### Surface reconstructions

Our analyses of folding, thickness and connectivity for the 6 primate sample used the b0 volumes from the diffusion-weighted MRI data. The data was upsampled to create high resolution surface reconstructions for all specimens except for the marmoset which was already scanned at very high resolution. The b0 image was used to manually segment the cerebrum using the same method as before: A first segmentation mask was obtained interactively using our tool Thresholdmann (https://neuroanatomy.github.io/thresholdmann, Heuer et al., 2024), which allows the creation of space-varying thresholds. These segmentations were then refined using BrainBox (https://brainbox.pasteur.fr, Heuer et al., 2016). BrainBox is a Web application providing several tools for collaborative manual segmentation of MRI data. The main task in BrainBox was to make sure that sulcal regions were open, and to correct additional segmentation and data artefacts. The masks generated by BrainBox were then used to produce surface meshes, using the implementation of the marching cubes algorithm provided by Scikit-Image (https://scikit-image.org, van der Walt et al. 2014). The meshes were then regularised using Graphite (https://github.com/BrunoLevy/GraphiteThree), which also automatically corrects a certain number of topological errors. Finally, the topology of the meshes was manually edited to obtain meshes of genus 2 (topologically spherical) using MeshSurgery (https://github.com/neuroanatomy/MeshSurgery).

#### Folding maps

We computed cortical folding maps for all brains using meshgeometry (https://github.com/neuroanatomy/meshgeometry). We used an algorithm which integrates mean curvature maps across different scales (Toro and Burnod, 2003). A map made from a single mean curvature map does not differentiate between deep and shallow folds. We computed 10 mean curvature maps at 10 different levels of Laplacian mesh smoothing, and added them. Deep folds stay present across all scales, which makes them appear more clearly in the integrated map (these are the maps shown in Fig. 4b in the main manuscript).

#### Comparisons

For pairwise comparison, the surface of one brain was resampled to the surface of the other brain based on the surface matching described above for establishing the homologies. The correlation between two brains was estimated as the average of the Pearson correlation between curvature maps resampled to the subdivided icosahedron (5 levels of subdivision) of each of the two brains.

#### Significance of cortical folding comparisons

The significance of the correlations was assessed by comparing the observed correlations to the ones obtained after random rotation of one of the spheres (spin test method, 1000 iterations, Vos de Wael et al., 2020). We applied hierarchical clustering on the correlation matrix for visualisation of the correlations (using the average for calculating the distance between newly formed clusters), Fig. S2.

#### Comparison of differences

We Fisher-transformed the curvature map correlations between species pairs and computed the differences between these transformed values, to obtain the difference in folding patterns between species.

To estimate the significance of those differences, we Fisher-transformed the correlations generated in the spin test permutations for each species pair. We computed the differences between those Fisher transformed correlations for each pair of species, and calculated the standard deviation which served as an estimate of the standard error as an insight on the significance of the folding similarities across species. To control for multiple comparisons, we applied false discovery rate (FDR, Benjamini and Hochberg,1995) correction to the p-values obtained from the standard error (Fig. S1).

#### Comparison of clusters

Intra-cluster Fisher-transformed correlations were on average 0.48 higher than inter-cluster Fisher-transformed correlations. In the spin test rotations (1000 iterations), the standard deviation of that difference was 0.02, leading to a highly significant p-value. Intra-cluster correlations were significantly higher than inter-cluster correlations (z_intra_-z_inter_=0.47, p≪1).

**Figure S2.**
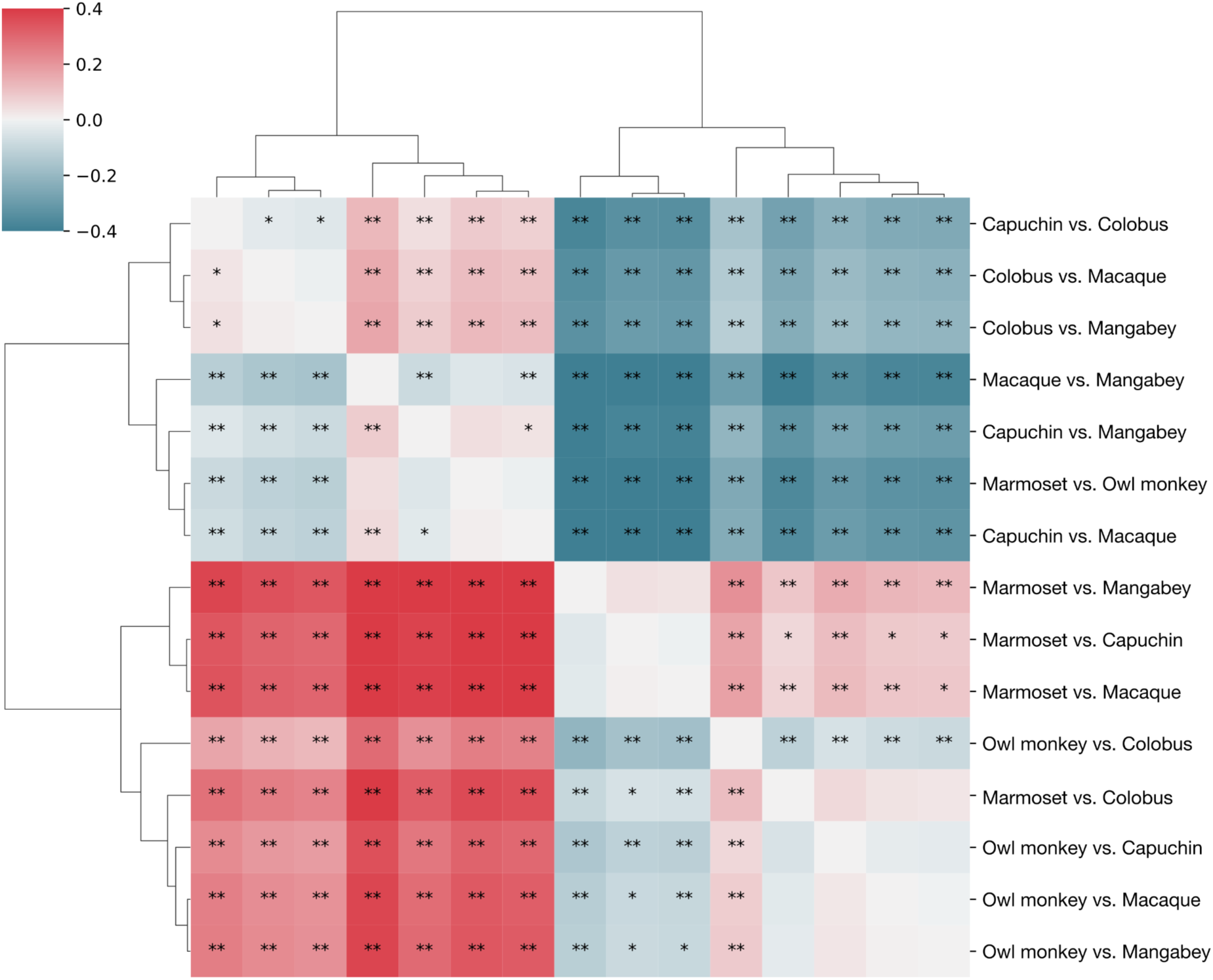
Significance of differences of cortical folding correlation between pairs of species. Differences of correlations marked with * and ** are significant with * FDR < 0.05; and ** FDR < 0.01. Species with similar brain volume are significantly more correlated, and species with different brain volumes are significantly less correlated.

### I.4 Cross-species comparison of cortical thickness

#### Cortical thickness computation

We automatically segmented the cortical ribbon across all species and derived cortical thickness maps.

First, each surface mesh was aligned with its MRI volume, and we computed a signed-distance function (SDF) for each volume. The value in each voxel in the SDF volume was 0 at the surface and negative inside of the surface. We then repeated the following procedure 100 times:

1. Parcellate the surface into 500 random parcels (using the Poisson disk method),
2. Assign each voxel in the volume to the closest surface parcel,
3. For all voxels within a parcel, compute the average grey-level value in the original MRI for all voxels closest to the surface, i.e., those whose SDF value was between 0 and −0.01,
4. Repeat the same in steps of 0.01 mm, to build a curve of grey-level changes as a function of SDF value (distance to the surface),
5. Smooth the grey-level versus distance curve by convolution with a Gaussian kernel and compute the point of steepest change: this is one local estimation of cortical thickness,
6. Assign this thickness value to all vertices in the parcel.

We created 100 parcellations per species (step 1), and then repeated steps 2 to 6 for all 100 different parcellations, and each time the vertices in the surface mesh belonged to a different parcel. To obtain a final estimation of cortical thickness for each vertex, we averaged all the 100 estimations coming from each parcellation.

The cortical thickness map in the surface mesh was then transformed into a volumetric segmentation. We computed the distance from each voxel to the closest point in the surface mesh (given in general by the barycentric coordinates of the triangle closest to the voxel). The thickness value for the voxel was taken as the weighted average of the cortical thickness values of the 3 vertices of this triangle. If the distance from the voxel to the surface mesh was smaller than the estimated cortical thickness, the voxel was marked as belonging to the cortex.

The automatic cortical segmentations were visually inspected and manually refined using BrainBox (Heuer et al., 2016). This refinement was then projected back to the cortical thickness map using a Euclidean distance transformation.

#### Validation

To assess the validity of our thickness estimation method, we compared the thickness map obtained for our rhesus macaque to the Yerkes macaque thickness map used by Hayashi et al. (2021), which represents the average of 19 macaques. The two methods provided very similar results (r=0.74, p≪1, Fig. S3).

#### Comparison of cortical thickness maps

For each species, we projected the brain surface mesh to a sphere (using the method described in Toro and Burnod (2003)) and resampled the thickness map to the subdivided icosahedron (5 subdivisions) of the sphere of each of the two brains. We then computed Pearson correlation between the thickness maps across species. Finally, we used the spin test method (1000 spins, Vos de Wael et al., 2020), as well as the sampling distribution and confidence intervals to assess the significance of the correlations (see Fig. S4).

#### Clustering

We performed a hierarchical clustering on the correlation matrix for visualisation of the correlations (using the average for calculating the distance between newly formed clusters). We computed the correlation of the thickness maps across species resampled to the surface sphere. For each pair of species this gave us 2 slightly different correlation values depending on which of the two species was used for computing the correlation. We averaged these two correlations (using a Fisher transform) to obtain a final symmetric matrix on which the clustering was performed.

**Figure S3.**
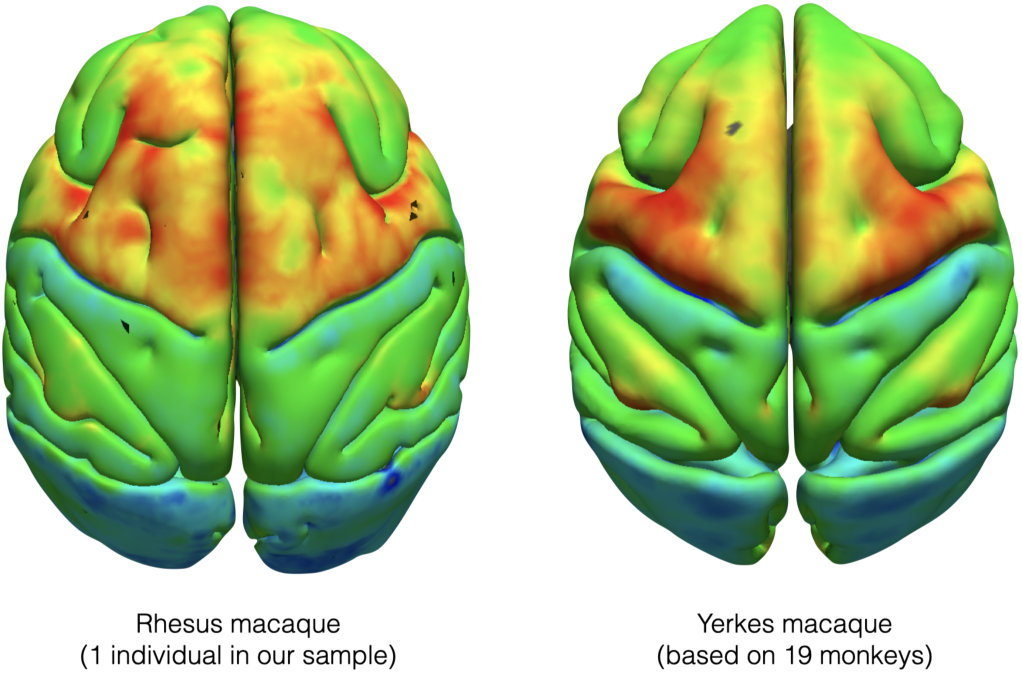
Validation of our cortical thickness estimation. Thickness map from the rhesus macaque in our sample compared to the Yerkes macaque thickness map from Hayashi et al. (2021).

#### Significance of cortical thickness comparisons

After computing the correlations of cortical thickness maps across the 6 species in our sample, we used the spin test permutation method (1000 spins) to evaluate if the observed similarities and differences between species’ cortical thickness maps were significant (Fig. S4). Species with similar brain volume are significantly more correlated, and species with different brain volumes are significantly less correlated.

#### Comparison of differences

Thickness map correlations between species pairs were Fisher-transformed and the differences between these transformed values were computed to quantify the difference in folding patterns between species.

To estimate the significance of these differences, we Fisher-transformed the correlations generated in the spin test permutations for each species pair. We then calculated the differences between the Fisher transformed correlations for each pair of species and used the standard deviation as an estimate of the standard error, providing insight into the significance of the thickness similarities across species. To control for multiple comparisons, false discovery rate (FDR, Benjamini and Hochberg,1995) correction was applied to the p-values obtained from the standard error (Fig. S4).

#### Comparison of clusters

Intra-cluster Fisher-transformed correlations were on average 0.35 higher than inter-cluster Fisher-transformed correlations. In the spin test rotations (1000 iterations), the standard deviation of that difference was 0.05, leading to a highly significant p-value. Intra-cluster correlations were significantly higher than inter-cluster correlations (z_intra_-z_inter_=0.34, p≪1).

**Figure S4.**
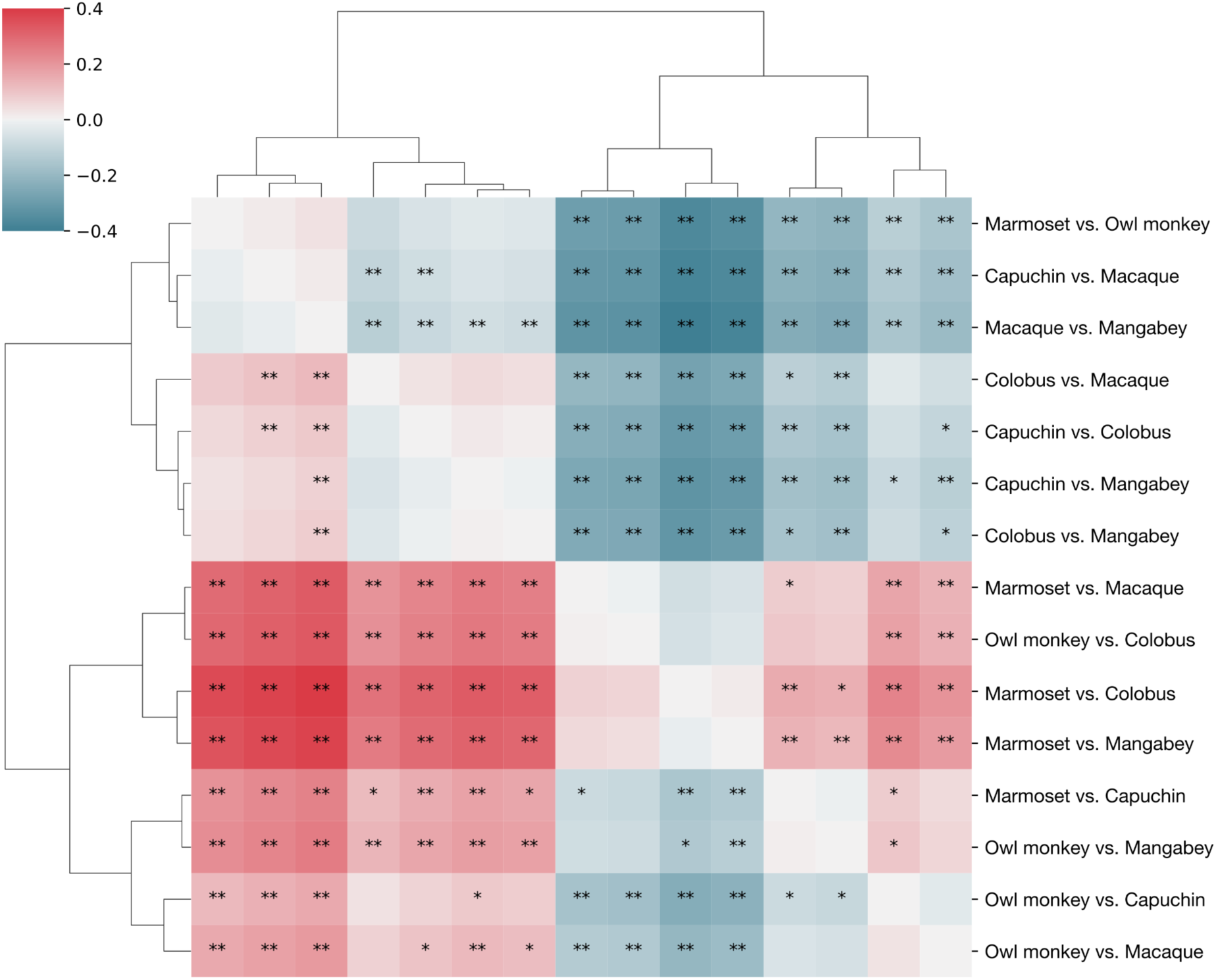
Significance of differences of cortical thickness correlation between pairs of species. Differences of correlations marked with * and ** are significant with * FDR < 0.05; and ** FDR < 0.01. Species with similar brain volume are significantly more correlated, and species with different brain volumes are significantly less correlated.

### I.5 Cross-species comparison of connectomes

We used a method based on random cortical parcellations to compare connectomes (illustrated in Fig. S5). The preliminary steps are (1) the surface morphing between all brain surfaces (Fig. S5a, b), (2) the computation of whole brain tractography (Fig. S5c, method described in the main Methods section).

#### Connectome computation

After this, we perform several iterations of the following algorithm (Fig. S5d):

1. We pick 1 of the 6 species.
2. We compute a random parcellation for this species.
3. The random parcellation is propagated to the other 5 species using the results of our surface morphing (section I.1).
4. For each species, we used Mrtrix (Tournier et al. 2019) to compute a connectome by counting the number of streamlines that go from parcel i to parcel j, for each pair of i, j, i≠j.
5. We pick each species 100 times, repeating steps 1 to 4, and use 100 random parcellations per species, leading to a total of 600 random parcellations per species, and 3600 connectomes (each parcellation is used for 6 connectomes, Fig. S5e left).
6. To account for a potential effect of the tracking, we computed tractograms using 3 different Fractional Anisotropy (FA) cutoff values: 0.02, 0.04, and 0.06.
7. To account for a potential effect of the number of parcels, we repeat all steps to obtain 6 sets of random parcellations with different amounts of parcels: 50, 100, 150, 200 (used in the main analyses), 250, and 300.

Fig. S5 shows an example of a connectome computed using this method.

#### Connectome comparison

For each set of 6 connectomes that correspond to a set of homologous parcellations, we used Spearman correlation to compute the similarity of the connectomes between pairs of species. We applied Fisher transformation to the correlation values to normalise their distribution. For each pair of species, we computed the mean Fisher transformed correlations across all connectomes.

We then applied the inverse Fisher transformation to obtain the final connectome correlation matrix for the 6 species. Hierarchical clustering was performed on the correlation matrix for visualisation (Fig. S5e right). The significance of the correlations between species pairs for each parcellation was estimated using a Mantel test with 100 permutations.

To account for a potential effect of the tracking, we repeated this procedure for 3 sets of connectomes computed with different FA cutoff values.

To account for a potential effect of the number of parcels, we repeated this procedure in the 6 sets of parcellations with different amounts of parcels.

**Figure S5.**
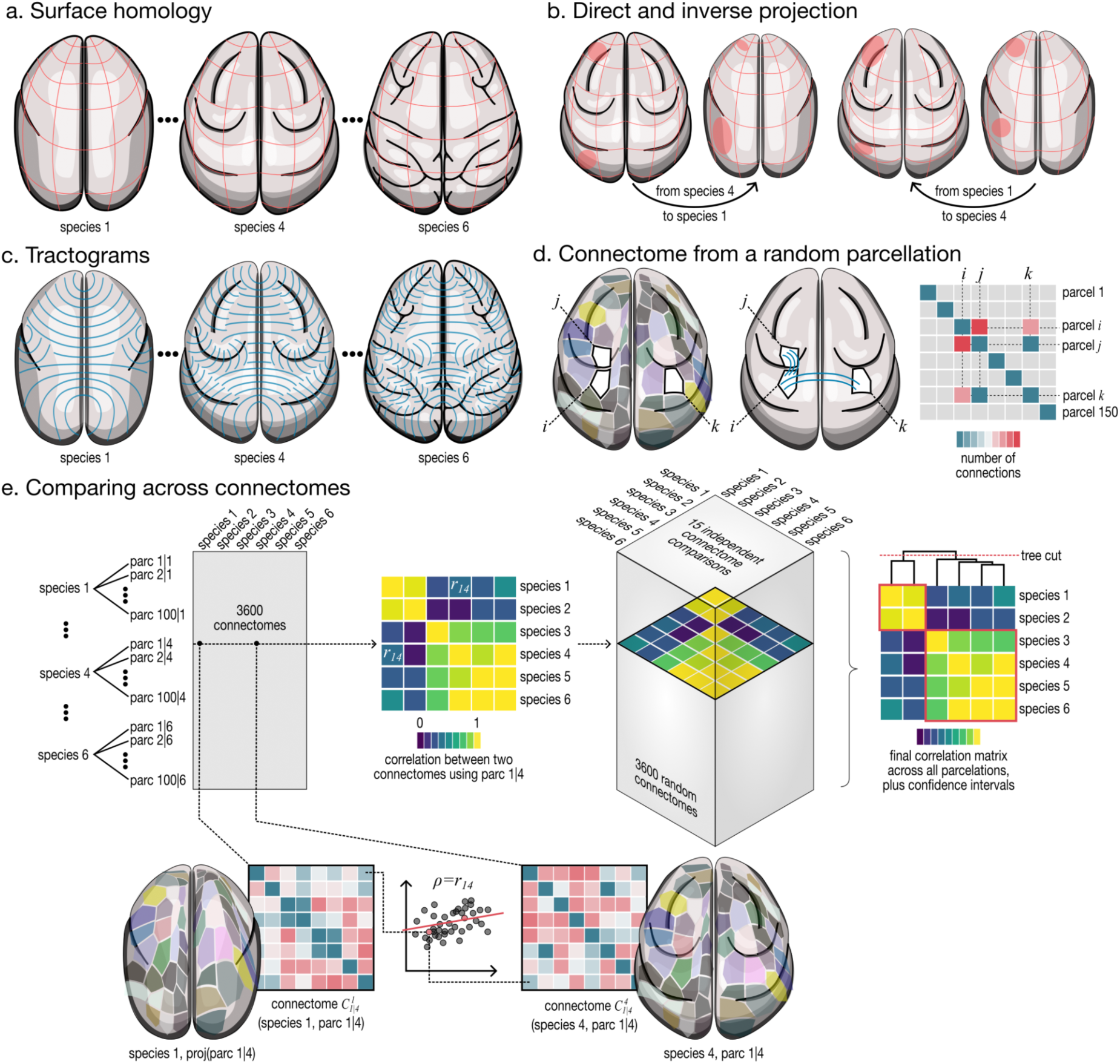
Connectome computation and comparison. The pre-requisites for the procedure are the computation of surface homologies (a), which enables direct and inverse projection of surface data (b); and the computation of whole-brain tractograms (c). Then, for the computation of one connectome, we start by computing a random parcellation of the surface. Based on this parcellation we compute a connectome by counting the number of fibres that link every parcel i and parcel j, with i≠j (d). To account for a potential effect of the species used for the parcellation, and for the geometry of one specific parcellation, we compute 100 parcellations for each species, which are then propagated to the other species, and we do that for each of the 6 species (e). The final similarity matrix is obtained by averaging across the similarity matrices of all those 3600 random connectomes. Nomenclature: parc i|j = parcellation number i computed on species j; proj(parc i|j) = projection of the parcellation parc i|j onto the given species; C^i_{j|k} = Connectome of species i (i=1,…, 6), computed using the parcellation parc j|k.

**Figure S6.**
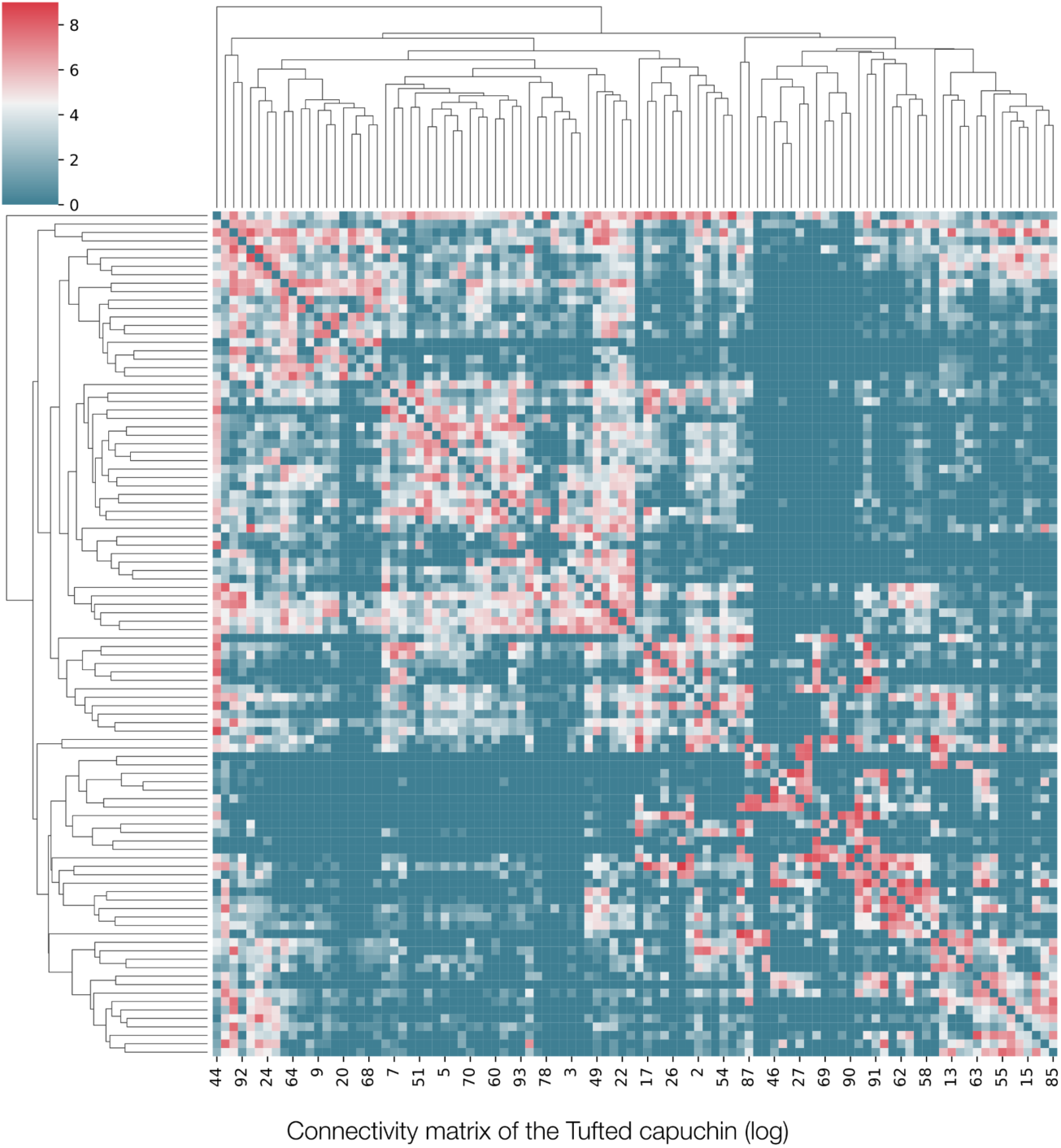
Example: One connectome of the capuchin monkey. Example connectome of one of the species (capuchin monkey) for one random parcellation with 100 parcels.

#### Comparison of differences

We Fisher-transformed the connectome correlations between species pairs, computed the differences between these transformed values, and averaged the differences across all parcellations to obtain the average difference in connectivity patterns between species.

To estimate the significance of those differences, we Fisher-transformed the correlations generated by the Mantel permutations for each species pair. We computed the differences between these Fisher transformed correlations for each pair of species across the Mantel iterations, and calculated the standard deviation of those differences. Finally, we averaged these values across all parcellations to obtain an estimate of the standard error as an insight on the significance of the connectome similarities across species. To control for multiple comparisons, we applied false discovery rate (FDR, Benjamini and Hochberg,1995) correction to the p-values obtained from the standard error (Fig. S7).

#### Comparison of clusters

Intra-cluster Fisher-transformed correlations were on average 0.15 higher than inter-cluster Fisher-transformed correlations. In the Mantel test permutations (100 times 600 permutations), the standard deviation of this difference was 0.013, leading to a significantly high p-value. Intra-cluster correlations were significantly higher than inter-cluster correlations (z_intra_-z_inter_=0.15, p≪1).

**Figure S7.**
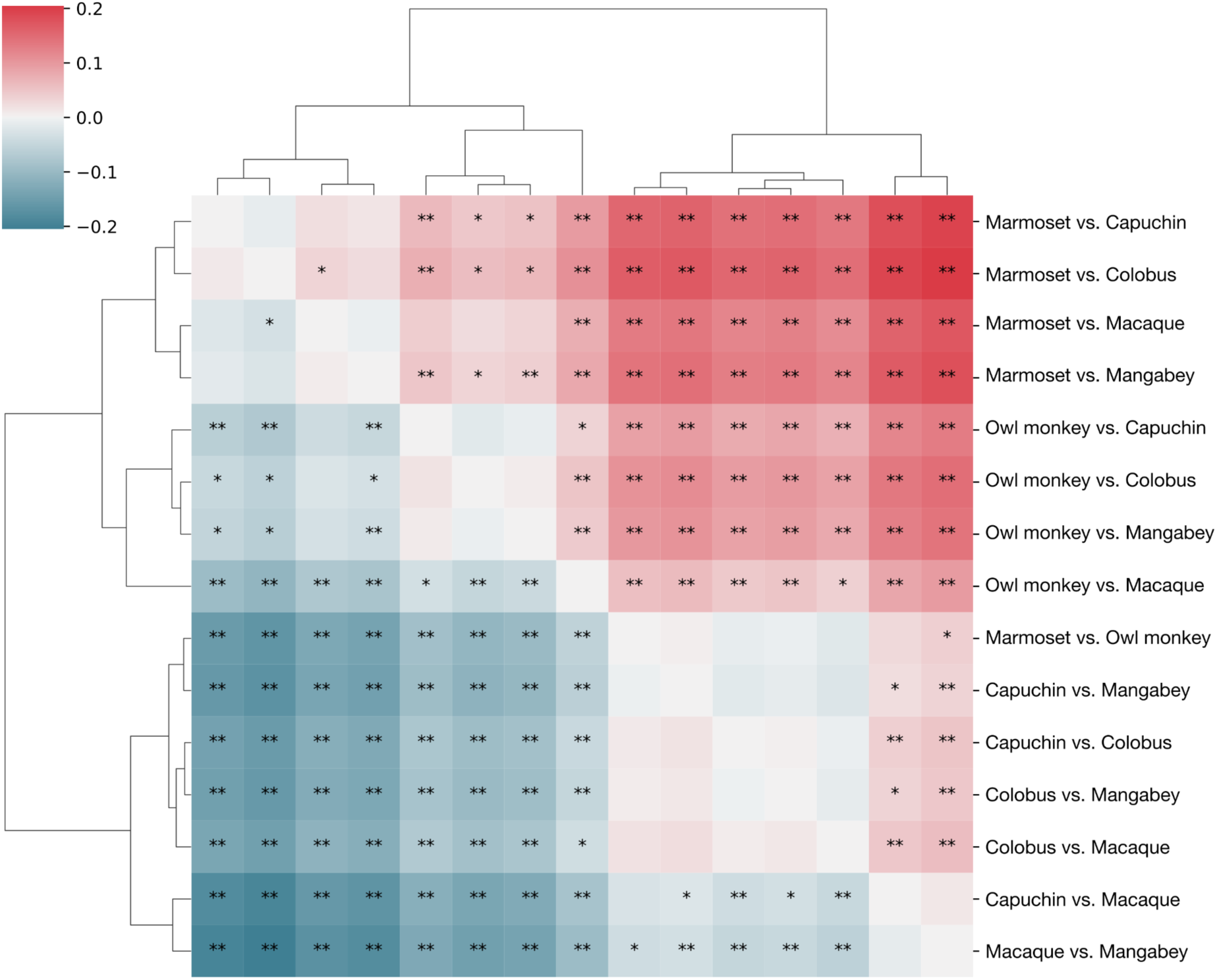
Significance of differences of cortico-cortical connectivity correlation between pairs of species. Differences of correlations marked with * and ** are significant with * FDR < 0.05; and ** FDR < 0.01. Species with similar brain volume are significantly more correlated, and species with different brain volumes are significantly less correlated.

#### Effect of different numbers of clusters on connectivity correlations

We computed the similarity of connectivity patterns across species for 50, 100, 150, 200, 250 and 300 parcels to test for a potential effect of parcel number on the observed correlations (Fig. S7). Our conclusions are robust to the number of parcels. The only effect we see is an overall decrease in all pairwise correlations with an increasing number of clusters, likely an effect of smoothing.

**Figure S8.**
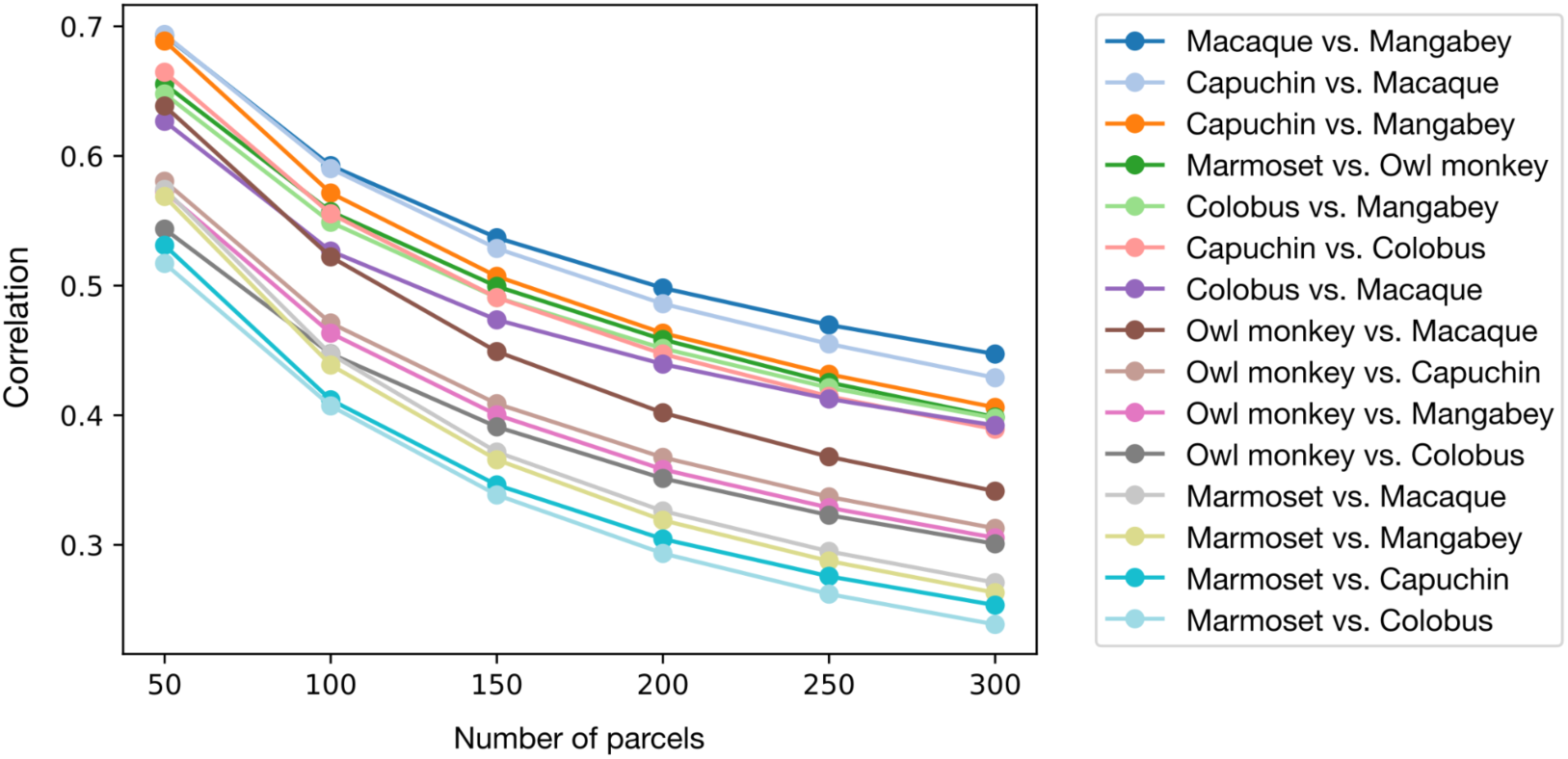
Effect of varying number of parcels on the species connectome comparisons. The similarity of connectivity patterns across species is shown for 50, 100, 150, 200, 250 and 300 parcels. Our conclusions are robust to the number of parcels. The only effect we see is an overall decrease in all pairwise correlations with an increasing number of clusters, likely an effect of smoothing.

#### Effect of different FA cutoff values on connectivity correlations

In order to track streamlines into the cortex, we tracked with a low FA value (see Guevara et al 2020) to terminate the fibres (FA=0.02). We used the cerebral masks for tracking.

Results from FA cutoff 0.02 (determined by visual inspection of the resulting connectome) were subsequently used in the main analyses reported in the paper. To test for a potential effect of the tracking cutoff value on the results, we performed the same analyses for cutoff values 0.04 and 0.06. The results and conclusion are robust across different cutoff values, showing the same pattern of high correlations between species with similar brain volume, and low correlations between species with different brain volume independent of phylogenetic relationship. Similar to the main analyses, we show here the results for 200 clusters. Figure S9 shows the correlation matrices for each cutoff value, and Figure S10 shows the mean correlation per species pair across the different cutoff values.

**Figure S9.**
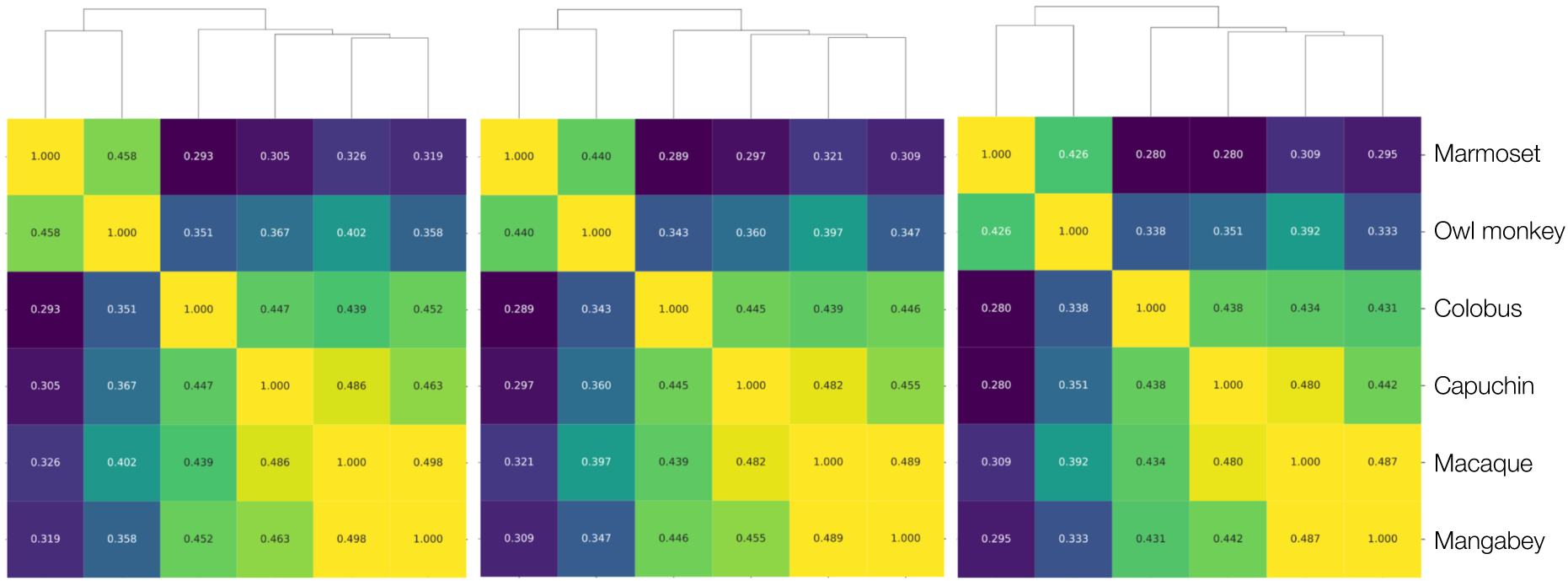
Correlation of connectivity patterns across species at different FA cutoff values. Left: 0.02 (reported in the main analyses), centre: 0.04, right: 0.06; here shown for 200-parcels parcellation. Results are robust across different cutoff values, showing that brains with similar brain volume have more similar connectivity patterns than brains of species with different brain volume independent of their phylogenetic relatedness.

**Figure S10.**
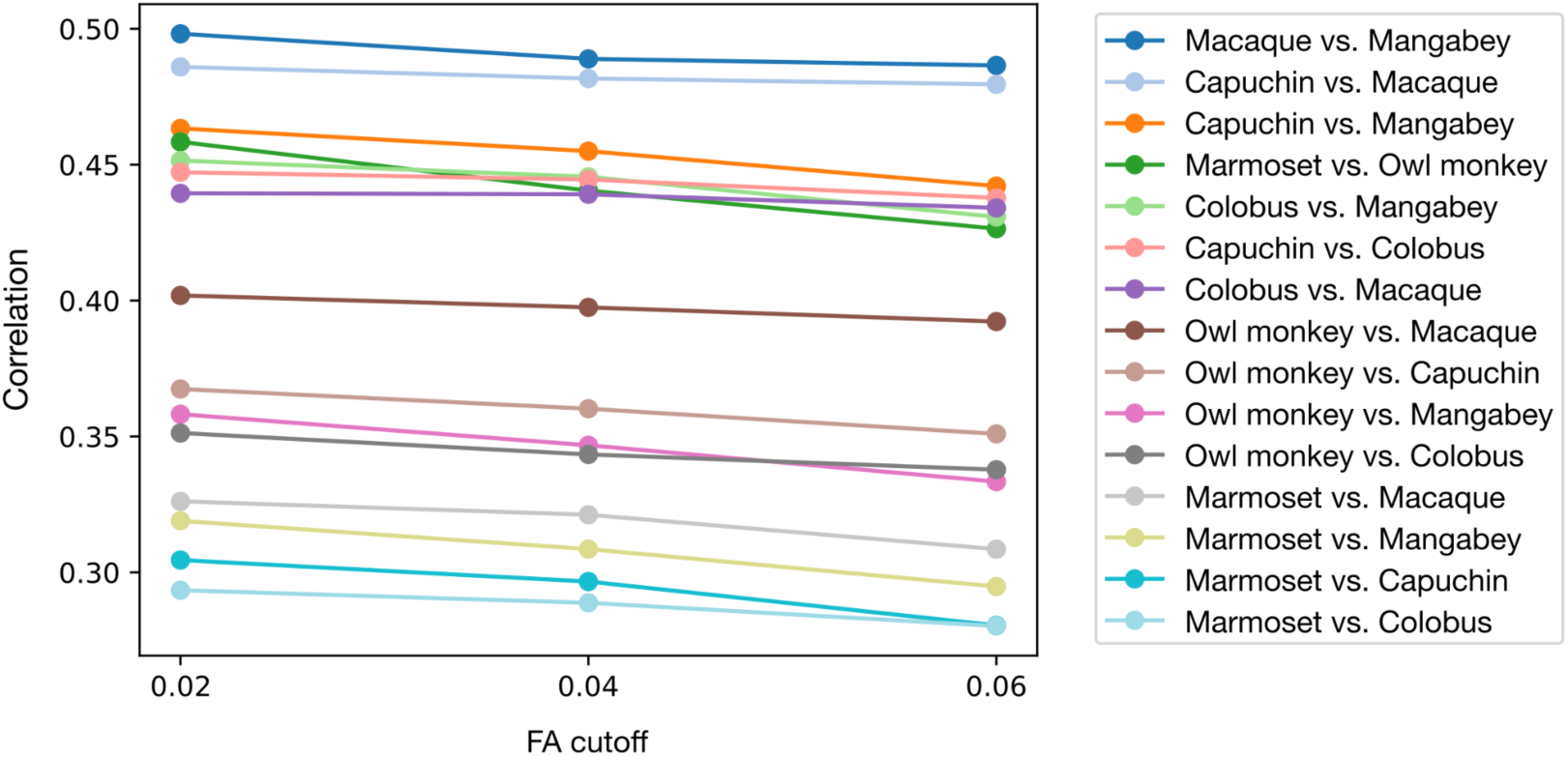
Mean correlation per species pair at different FA cutoff values. Results are robust across different cutoff values, showing that brains with similar brain volume have more similar connectivity patterns than brains of species with different brain volume independent of their phylogenetic relatedness. Left: FA cutoff value 0.02 (reported in the main analyses), centre: 0.04, right: 0.06; here shown for 200-parcels parcellation.

### I.6 Cross-species comparison of behaviour

To check for a potential effect of the phylogenetic tree on our analyses results, we obtained phylogenetic tree data from an additional source, the 10k trees website (Arnold et al. 2010, https://10ktrees.nunn-lab.org/Primates/downloadTrees.php), which provides a Bayesian inference of primate phylogeny. We downloaded the consensus tree, and performed the same analyses as reported in the main paper based on this phylogenetic tree.

Our conclusions are robust to the choice of the phylogenetic tree. We obtained the same correlation values between the behavioural and phylogenetic distance matrices for both phylogenetic trees (see Figure S11).

**Figure S11.**
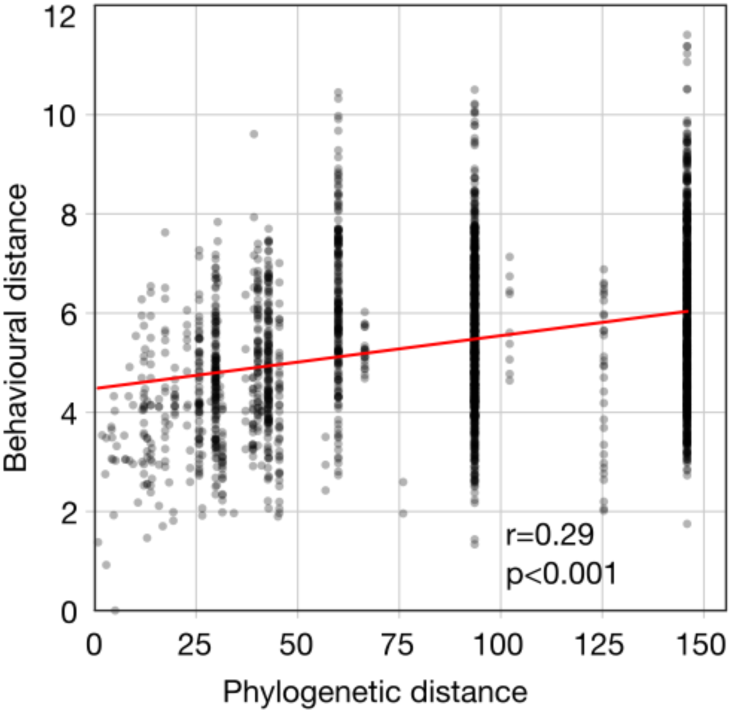
Behavioural and phylogenetic similarity across species. The correlation between phylogenetic similarity and behavioural similarity based on the 10k trees phylogenetic tree (Arnold et al. 2010) was the same as when performed on the phylogenetic tree from Dos Reis et al. (2018): very low, with just a slightly tighter 95% confidence interval (r=0.29, p<0.001, 95% CI=[0.19, 0.39].

## II. Videos

### II.1 Evolutionary expansion trajectory across 54 species

**Video 1. Evolutionary expansion trajectory across 54 primate species.** The video shows a neocortical expansion trajectory across 54 primate species, based purely on brain size, starting from small lemurs and going towards great apes. The dot in the phylogenetic tree indicates the last common ancestor of 2 adjacent brains in the trajectory. Despite jumping to far apart branches of the phylogenetic tree, the resulting trajectory is strikingly continuous.

### II.2 Surface matching between pairs of species

**Video 2. Morphing between 2 species which are phylogenetically far.** Surface morphing between a capuchin brain (New World monkey) and a macaque brain (Old World monkey) shows that the folding pattern of capuchins closely matches that of the Old World monkey, despite having evolved largely independently. Although these species are far in the phylogenetic tree, morphing between them is extremely smooth and shows almost no deformation due to the high similarity of the brain geometry.

## III. Supplementary Tables

### III.1 Evolutionary model selection

#### III.1.1 Model selection for 70 primate species

**Table S1.** Phylogenetic model selection. Different models of phenotypic character evolution were fitted to the data and ranked by their Akaike information criterion corrected for small sample sizes (AICc). The smallest value indicates the best fit.

| Model | AICc |
| --- | --- |
| Brownian Motion | -550.36 |
| Early Burst | -548.03 |
| Ornstein Uhlenbeck | -535.44 |

#### III.1.2 Model selection for 49 New World monkey species

**Table S2.**
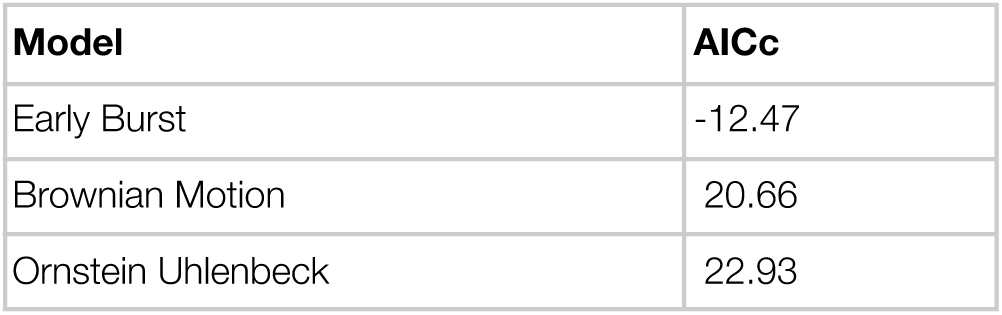
Phylogenetic model selection. Different models of phenotypic character evolution were fitted to the data and ranked by their Akaike information criterion corrected for small sample sizes (AICc). The smallest value indicates the best fit.

### III.2 Overview of the collected behavioural and neuroanatomical variables

**Table S3.** Behavioural and neuroanatomical data. We collected data on primate behaviour from the scientific literature as well as online databases. We also computed detailed neuroanatomical measurements for 70 primate species based on our surface reconstructions, and included endocranial and brain volume data from the literature.

|  |
| --- |
| **Behavioural data (20)** |
| Group size, Tool use, Innovation, Extractive foraging, Innovation frequency, Tool use frequency, Extractive foraging frequency, Social learning frequency, Tool manufacturing, Motor complexity, Knowledge score, Manipulation complexity, Diet (folivore, frugivore, omnivore), Diet composition (percentage of fruits, insects, leaves), Home range size, Activity period (diurnal, nocturnal, cathemeral), Mating system (harem polygyny, monogamy, polyandry, polygynandry, spatial polygyny), Social system (pair, polygynandry, polygyny, solitary), Habitat/Locomotion (arboreal/terrestrial), and Social learning. |
| **Neuroanatomical data (5)** |
| Endocranial volume, Brain volume, Cerebral volume, Surface area, Absolute gyrification index |

### III.3 List of species in the cerebral surface analyses

**Table S4.**
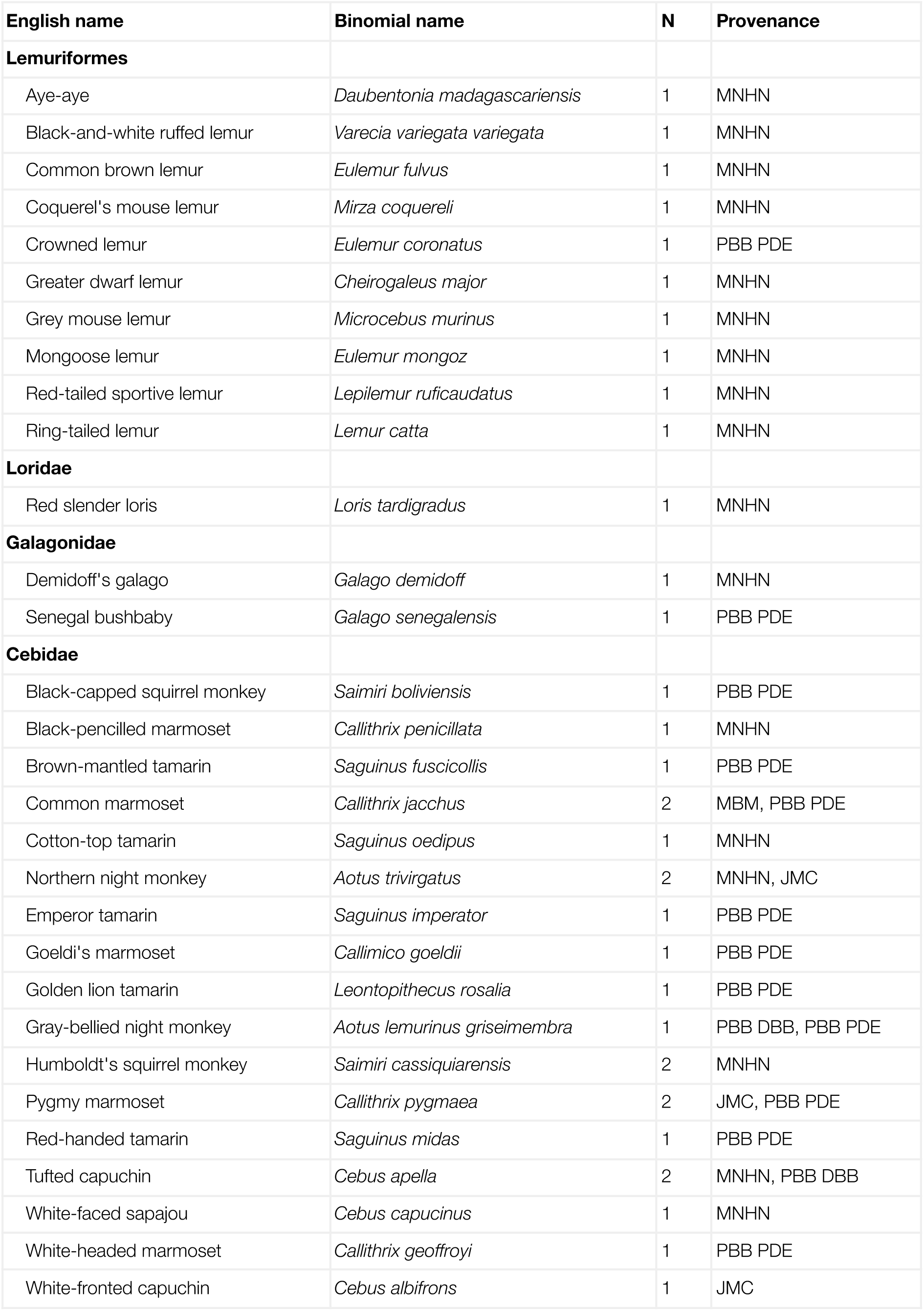
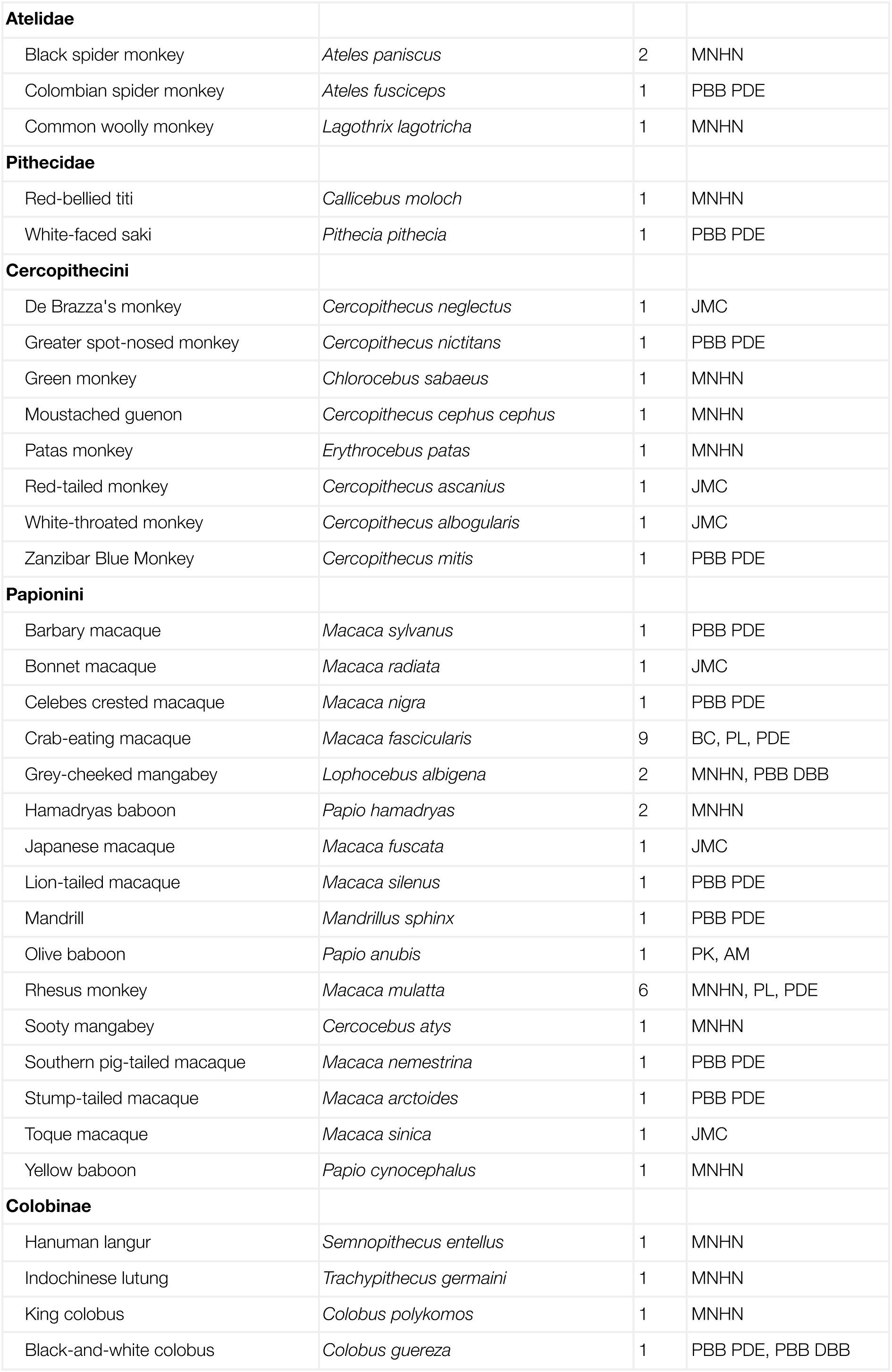
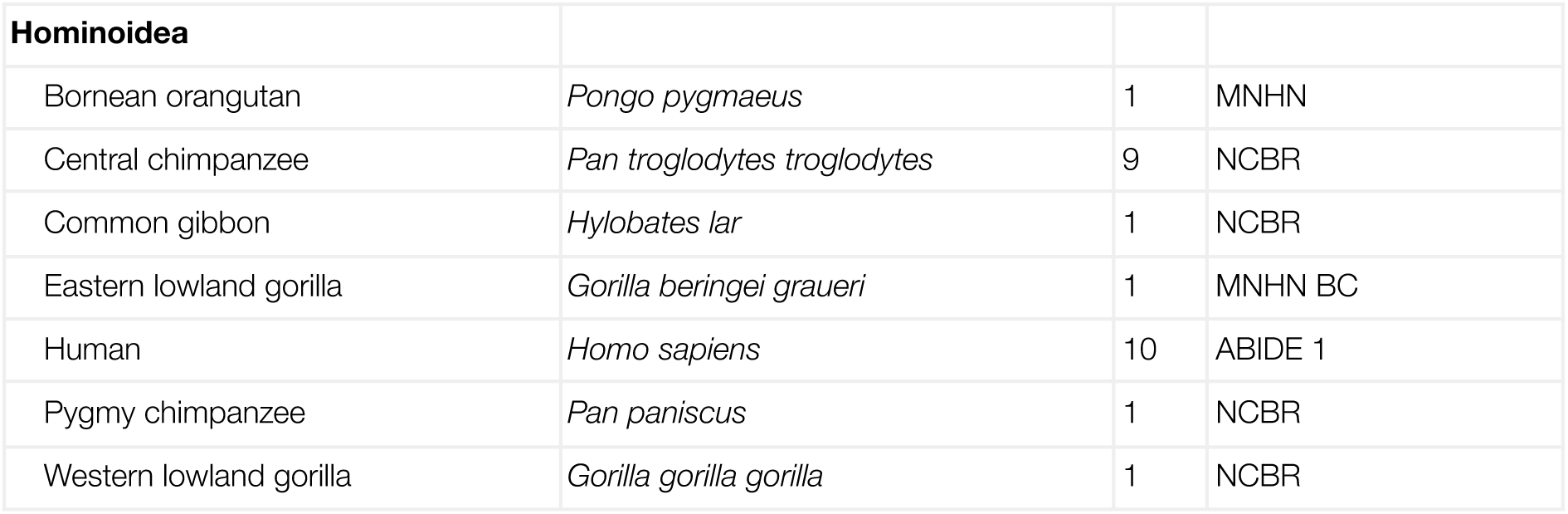
List of species included in the MRI dataset. ABIDE 1: Autism Brain Imaging Data Exchange 1. BC: Brain Catalogue. MNHN: Muséum Nationale d’Histoire Naturelle de Paris. NCBR: National Chimpanzee Brain Resource. PL: Pruszynski Lab. PBB: Primate Brain Bank. PDE: PRIMate Data Exchange (PRIME-DE). MBM: Marmoset Brain Mapping project. JMC: Japan Monkey Centre. DBB: Digital Brain Bank. AM: Adrien Meguerditchian. PK: Peter Kochunov.

III.3 List of species in the cerebral surface analyses.
| English name | Binomial name | N | Provenance |
| --- | --- | --- | --- |
| **Lemuriformes** |  |  |  |
| Aye-aye | *Daubentonia madagascariensis* | 1 | MNHN |
| Black-and-white ruffed lemur | *Varecia variegata variegata* | 1 | MNHN |
| Common brown lemur | *Eulemur fulvus* | 1 | MNHN |
| Coquerel's mouse lemur | *Mirza coquereli* | 1 | MNHN |
| Crowned lemur | *Eulemur coronatus* | 1 | PBB PDE |
| Greater dwarf lemur | *Cheirogaleus major* | 1 | MNHN |
| Grey mouse lemur | *Microcebus murinus* | 1 | MNHN |
| Mongoose lemur | *Eulemur mongoz* | 1 | MNHN |
| Red-tailed sportive lemur | *Lepilemur ruficaudatus* | 1 | MNHN |
| Ring-tailed lemur | *Lemur catta* | 1 | MNHN |
| **Loridae** |  |  |  |
| Red slender loris | *Loris tardigradus* | 1 | MNHN |
| **Galagonidae** |  |  |  |
| Demidoff's galago | *Galago demidoff* | 1 | MNHN |
| Senegal bushbaby | *Galago senegalensis* | 1 | PBB PDE |
| **Cebidae** |  |  |  |
| Black-capped squirrel monkey | *Saimiri boliviensis* | 1 | PBB PDE |
| Black-pencilled marmoset | *Callithrix penicillata* | 1 | MNHN |
| Brown-mantled tamarin | *Saguinus fuscicollis* | 1 | PBB PDE |
| Common marmoset | *Callithrix jacchus* | 2 | MBM, PBB PDE |
| Cotton-top tamarin | *Saguinus oedipus* | 1 | MNHN |
| Northern night monkey | *Aotus trivirgatus* | 2 | MNHN, JMC |
| Emperor tamarin | *Saguinus imperator* | 1 | PBB PDE |
| Goeldi's marmoset | *Callimico goeldii* | 1 | PBB PDE |
| Golden lion tamarin | *Leontopithecus rosalia* | 1 | PBB PDE |
| Gray-bellied night monkey | *Aotus lemurinus griseimembra* | 1 | PBB DBB, PBB PDE |
| Humboldt's squirrel monkey | *Saimiri cassiquiarensis* | 2 | MNHN |
| Pygmy marmoset | *Callithrix pygmaea* | 2 | JMC, PBB PDE |
| Red-handed tamarin | *Saguinus midas* | 1 | PBB PDE |
| Tufted capuchin | *Cebus apella* | 2 | MNHN, PBB DBB |
| White-faced sapajou | *Cebus capucinus* | 1 | MNHN |
| White-headed marmoset | *Callithrix geoffroyi* | 1 | PBB PDE |
| White-fronted capuchin | *Cebus albifrons* | 1 | JMC |
| **Atelidae** |  |  |  |
| Black spider monkey | *Ateles paniscus* | 2 | MNHN |
| Colombian spider monkey | *Ateles fusciceps* | 1 | PBB PDE |
| Common woolly monkey | *Lagothrix lagotricha* | 1 | MNHN |
| **Pitheciidae** |  |  |  |
| Red-bellied titi | *Callicebus moloch* | 1 | MNHN |
| White-faced saki | *Pithecia pithecia* | 1 | PBB PDE |
| **Cercopithecini** |  |  |  |
| De Brazza's monkey | *Cercopithecus neglectus* | 1 | JMC |
| Greater spot-nosed monkey | *Cercopithecus nictitans* | 1 | PBB PDE |
| Green monkey | *Chlorocebus sabaeus* | 1 | MNHN |
| Moustached guenon | *Cercopithecus cephus cephus* | 1 | MNHN |
| Patas monkey | *Erythrocebus patas* | 1 | MNHN |
| Red-tailed monkey | *Cercopithecus ascanius* | 1 | JMC |
| White-throated monkey | *Cercopithecus albogularis* | 1 | JMC |
| Zanzibar Blue Monkey | *Cercopithecus mitis* | 1 | PBB PDE |
| **Papionini** |  |  |  |
| Barbary macaque | *Macaca sylvanus* | 1 | PBB PDE |
| Bonnet macaque | *Macaca radiata* | 1 | JMC |
| Celebes crested macaque | *Macaca nigra* | 1 | PBB PDE |
| Crab-eating macaque | *Macaca fascicularis* | 9 | BC, PL, PDE |
| Grey-cheeked mangabey | *Lophocebus albigena* | 2 | MNHN, PBB DBB |
| Hamadryas baboon | *Papio hamadryas* | 2 | MNHN |
| Japanese macaque | *Macaca fuscata* | 1 | JMC |
| Lion-tailed macaque | *Macaca silenus* | 1 | PBB PDE |
| Mandrill | *Mandrillus sphinx* | 1 | PBB PDE |
| Olive baboon | *Papio anubis* | 1 | PK, AM |
| Rhesus monkey | *Macaca mulatta* | 6 | MNHN, PL, PDE |
| Sooty mangabey | *Cercocebus atys* | 1 | MNHN |
| Southern pig-tailed macaque | *Macaca nemestrina* | 1 | PBB PDE |
| Stump-tailed macaque | *Macaca arctoides* | 1 | PBB PDE |
| Toque macaque | *Macaca sinica* | 1 | JMC |
| Yellow baboon | *Papio cynocephalus* | 1 | MNHN |
| **Colobinae** |  |  |  |
| Hanuman langur | *Semnopithecus entellus* | 1 | MNHN |
| Indochinese lutung | *Trachypithecus germaini* | 1 | MNHN |
| King colobus | *Colobus polykomos* | 1 | MNHN |
| Black-and-white colobus | *Colobus guereza* | 1 | PBB PDE, PBB DBB |

**Table S4. List of species included in the MRI dataset.**
| **Hominoidea** |  |  |  |
| --- | --- | --- | --- |
| Bornean orangutan | *Pongo pygmaeus* | 1 | MNHN |
| Central chimpanzee | *Pan troglodytes troglodytes* | 9 | NCBR |
| Common gibbon | *Hylobates lar* | 1 | NCBR |
| Eastern lowland gorilla | *Gorilla beringei graueri* | 1 | MNHN BC |
| Human | *Homo sapiens* | 10 | ABIDE 1 |
| Pygmy chimpanzee | *Pan paniscus* | 1 | NCBR |
| Western lowland gorilla | *Gorilla gorilla gorilla* | 1 | NCBR |

### III.4 List of species in the endocast analyses

**Table S5.** List of species included in the endocast dataset. Endocasts of 179 specimens from 49 New World monkey species from Aristide et al. (2016).

| English name | Binomial name |
| --- | --- |
| **Atelidae** |  |
| Red-handed howler | *Alouatta belzebul* |
| Black howler | *Alouatta caraya* |
| Brown howler | *Alouatta guariba* |
| Guyanese red howler | *Alouatta macconnelli* |
| Mantled howler | *Alouatta palliata* |
| Yucatán black howler | *Alouatta pigra* |
| Colombian red howler | *Alouatta seniculus* |
| Peruvian spider monkey | *Ateles chamek* |
| Black-handed spider monkey | *Ateles geoffroyi* |
| White-cheeked spider monkey | *Ateles marginatus* |
| Red-faced spider monkey | *Ateles paniscus* |
| Southern muriqui | *Brachyteles arachnoides* |
| Northern muriqui | *Brachyteles hypoxanthus* |
| Common woolly monkey | *Lagothrix lagotricha* |
| **Cebidae** |  |
| Azara's night monkey | *Aotus azarae* |
| Black-headed night monkey | *Aotus nigriceps* |
| Spix's night monkey | *Aotus vociferans* |
| Goeldi's marmoset | *Callimico goeldii* |
| Buffy-tufted marmoset | *Callithrix aurita* |
| Common marmoset | *Callithrix jacchus* |
| Wied's marmoset | *Callithrix kuhlii* |
| Black-tufted marmoset | *Callithrix penicillata* |
| Pygmy marmoset | *Cebuella pygmaea* |
| White-fronted capuchin | *Cebus albifrons* |
| Tufted capuchin | *Cebus apella* |
| Colombian white-faced capuchin | *Cebus capucinus* |
| Crested capuchin | *Cebus robustus* |
| Golden-headed lion tamarin | *Leontopithecus chrysomelas* |
| Golden lion tamarin | *Leontopithecus rosalia* |
| Silvery marmoset | *Mico argentatus* |
| Gold-and-white marmoset | *Mico chrysoleucus* |
| Santarem marmoset | *Mico humeralifer* |
| Pied tamarin | *Saguinus bicolor* |
| Brown-mantled tamarin | *Saguinus fuscicollis* |
| Golden-handed tamarin | *Saguinus midas* |
| Moustached tamarin | *Saguinus mystax* |
| Black-capped squirrel monkey | *Saimiri boliviensis* |
| Common squirrel monkey | *Saimiri sciureus* |
| **Pitheciidae** |  |
| Bald uakari | *Cacajao calvus* |
| Golden-backed uakari | *Cacajao melanocephalus* |
| Brown titi monkey | *Callicebus brunneus* |
| White-eared titi monkey | *Callicebus donacophilus* |
| Red-bellied titi monkey | *Callicebus moloch* |
| Atlantic titi monkey | *Callicebus personatus* |
| Red-backed bearded saki | *Chiropotes chiropotes* |
| Black bearded saki | *Chiropotes satanas* |
| Gray's bald-faced saki | *Pithecia irrorata* |
| Monk saki | *Pithecia monachus* |
| White-faced saki | *Pithecia pithecia* |

## Notes

### Competing Interest Statement

The authors have declared no competing interest.

